# Selective advantage of mutant stem cells in clonal hematopoiesis occurs by attenuating the deleterious effects of inflammation and aging

**DOI:** 10.1101/2023.09.12.557322

**Authors:** Niels Asger Jakobsen, Sven Turkalj, Andy G. X. Zeng, Bilyana Stoilova, Marlen Metzner, Murtaza S. Nagree, Sayyam Shah, Rachel Moore, Batchimeg Usukhbayar, Mirian Angulo Salazar, Grigore-Aristide Gafencu, Alison Kennedy, Simon Newman, Benjamin J. L. Kendrick, Adrian H. Taylor, Rasheed Afinowi-Luitz, Roger Gundle, Bridget Watkins, Kim Wheway, Debra Beazley, Alexander Murison, Alicia G. Aguilar-Navarro, Eugenia Flores-Figueroa, Stephanie G. Dakin, Andrew J. Carr, Claus Nerlov, John E. Dick, Stephanie Z. Xie, Paresh Vyas

**Author notes:** These authors contributed equally (Niels Asger Jakobsen, Sven Turkalj, Andy G. X. Zeng). Senior author (John E. Dick, Stephanie Z. Xie, Paresh Vyas). Lead contact author: Paresh Vyas.

## Abstract

Clonal hematopoiesis (CH) arises when hematopoietic stem cells (HSC) acquire mutations in genes, including *DNMT3A* and *TET2*, conferring a competitive advantage through a mechanism that remains unclear. To gain insight into how CH mutations enable gradual clonal expansion, we used single-cell multi-omics with high-fidelity genotyping on CH bone marrow samples. Most of the selective advantage of mutant cells occurs within HSCs. *DNMT3A* and *TET2*-mutant clones expand further in early progenitors, while *TET2* mutations accelerate myeloid maturation in a dose-dependent manner. Unexpectedly, both mutant and non-mutant HSCs from CH samples are enriched for inflammatory and aging transcriptomic signatures, compared to HSC from non-CH samples, revealing a non-cell autonomous mechanism. However, *DNMT3A* and *TET2*-mutant HSCs have an attenuated inflammatory response relative to wild-type HSCs within the same sample. Our data support a model whereby CH clones are gradually selected because they are more resistant to the deleterious impact of inflammation and aging.

## Introduction

Aging tissues accumulate somatic mutations. If these occur in long-lived tissue stem cells, they provide a substrate for clonal selection and subsequent clonal expansion, resulting in somatic mosaicism.^1,2^ Somatic mosaicism in phenotypically normal human tissues was first described in blood and termed clonal hematopoiesis (CH).^3–9^ Subsequently, similar findings were documented in multiple other tissues.^10–19^ CH becomes common with aging and is associated with a 12-fold increased risk of myeloid malignancy, cardiovascular disease, and other adverse outcomes.^7–9,20^ A key unresolved question in the field is, what are the mechanisms that lead to selection of mutant clones? For a selected clone, increasing clone size has been shown to contribute the risk of myeloid malignancy,^21,22^ cardiovascular disease,^20,23^ and diseases associated with human aging.^24,25^ Though it is likely that multiple mechanisms contribute both positively and negatively to the rate of expansion of mutant clones,^26^ mechanistic understanding of the biological principles of expansion remain unclear.

Interestingly, ∼70% of CH cases are associated with mutations in just two genes, *DNMT3A* and *TET2*.^7,8,21,22,27^ DNMT3A, a *de novo* DNA methyltransferase, catalyzes the conversion of cytosine to 5-methylcytosine (5mC), usually in CpG dinucleotides.^28^ TET2 is a dioxygenase that catalyzes the conversion of 5-methylcytosine to 5-hydroxymethylcytosine (5hmC) and other oxidized derivatives.^29,30^ This reaction is the first step in DNA demethylation, although 5hmC can also act as a stable epigenetic mark with a regulatory role.^31,32^ Both proteins have also been associated with other roles, for example RNA splicing,^33^ regulation of RNA stability,^34–37^ and recruitment of other epigenetic regulators.^37–40^ In CH, *DNMT3A* mutations are predominantly heterozygous, scattered throughout the three functional domains and predicted to cause loss of function (LoF). In contrast, ∼60% of *DNMT3A* mutations in acute myeloid leukemia (AML) affect the R882 hotspot residue in the methyltransferase domain. *TET2* mutations are either missense or truncating variants distributed across the coding region and are predicted to inhibit or abolish the enzyme’s catalytic activity.

In mice, genetic mutants of *Dnmt3a* and *Tet2* confer clonal advantage to HSCs and alter their differentiation. Adult *Dnmt3a^-/-^* HSCs outcompete wild-type (WT) HSCs in secondary, and subsequent, competitive transplants.^41^ Similarly, *Tet2^-/-^* HSCs have a competitive advantage over WT HSCs in transplantation assays,^42–45^ although this advantage is most marked in early serial transplants.^46^ *Tet2^+/-^* HSCs have a similar competitive advantage but with longer latency.^44^ The competitive advantage of *Dnmt3a*^-/-^ HSCs has been attributed to increased RNA expression of selected multipotency and self-renewal genes, which correlates with hypomethylation of their promoters and gene bodies.^41^ Re-expression of DNMT3A in *Dnmt3a*^-/-^ HSCs partially restores methylation at specific genes and reduces HSC frequency.^41^ Notably, clonal advantage of *Dnmt3a^-/-^* HSCs is increased when mice are exposed to infection and inflammation, and abrogated when receptors for interferon-γ and TNFα are deleted.^47,48^ Aging and inflammation also expands both immunophenotypic and functional *Tet2*^−/−^ and *Tet2*^+/−^ HSCs.^49–51^ This is mimicked by IL-6 and IL-1 exposure and partially abrogated by blocking their cognate receptors.^49–51^ HSC differentiation is also altered in mouse models of CH mutations. Acute bi-allelic deletion of *Dnmt3a* modestly increases transcriptionally defined megakaryocyte-erythroid cells, with a commensurate decrease in myelomonocytic cells.^52^ Altered differentiation of *Dnmt3a*^−/−^ cells may be caused by decreased methylation of binding motifs for erythroid versus myeloid transcription factors (TFs), due to differences in motif CpG content.^52^ *Tet2*^−/−^ cells have a myeloid bias with expansion of GMP and reduction of megakaryocyte erythroid (MEP) and lymphoid progenitors.^52^ With age, some *Tet2*^−/−^ mice develop chronic myelomonocytic leukemia (CMML).^43–45^

However, the murine studies may not capture all the complexities associated with human CH. In humans, CH arises when a single cell acquires a mutation that confers a selective advantage, leading to gradual clonal expansion over time. Indeed, recent studies using population genetic modelling, single-cell phylogenetic analysis, and longitudinal sampling have estimated that *DNMT3A* and *TET2* mutant CH clones expand by about 5–20% per year and are acquired decades before they reach a substantial clone size.^53–55^ In contrast, mice are kept in controlled environments, have short lifespans and murine studies often assay hemopoiesis after transplantation or introduction of a mutant allele in all blood cells, rather than study HSC competition dynamics in native hematopoiesis with introduction of subclonal mutations.

In humans, detailed single-cell analyses have been performed in one genetic CH subtype (*DNMT3A* R882^+/-^) from patients treated for myeloma^56^ and in cord blood hematopoietic stem and progenitor cells (HSPCs) where both *TET2* alleles were experimentally deleted.^57^ In both studies, differentiation defects were observed, with engineered *TET2*^-/-^ cells having a competitive advantage *in vivo* in immunodeficient mice. However, it is unclear from these studies how *DNMT3A* and *TET2* mutations confer a gradual clonal advantage to mutant HSC in age-associated CH. To address this question, we implemented an optimized single-cell multi-omics platform, combining high-fidelity genotyping with high-quality transcriptional profiling, to analyze mutant HSPCs separately from non-mutant HSPCs from the same individuals, enabling us to dissect the cell-intrinsic and cell-extrinsic consequences of CH mutations in human BM.

## Results

### Bone marrow sampling of HSPCs from individuals with age-related clonal hematopoiesis without prior malignancy

As our goal was to study CH-mutant HSPCs and compare their properties to non-mutant HSPCs from the same individual, we needed to obtain bone marrow samples from individuals with CH, but without the confounding effects of co-existing malignancy or known inflammatory conditions. We collected samples from 195 individuals at various ages with normal blood counts undergoing elective total hip replacement surgery for osteoarthritis. To study steady-state CH, we excluded subjects with prior or current hematological cancer, inflammatory arthritis, or systemic steroid use (clinical characteristics, Table S1). We screened samples for CH by targeted re-sequencing of bone marrow mononuclear cell (BM MNC) DNA using a 347 kb panel covering 97 genes to a mean depth of 822X (Figure 1A, Figure S1A, Table S2). 57 of 195 individuals (29.2%) had CH with somatic driver mutation(s) at a variant allele frequency (VAF) ≥ 0.02, and an additional 28 individuals (14.3%) had CH with mutation(s) at a VAF of 0.01–0.02 (Figure 1B-C, Table S2). Consistent with prior studies, 69% of CH cases had mutations in *DNMT3A* and *TET2* predicted to cause loss-of-function (Figure S1B). The median VAF of all mutations detected in the cohort was 0.022 (Figures 1D). The frequency of secondary mutations increased with age and two or more mutations were observed in 45% of CH cases over the age of 80 (Figure 1E).

**Figure 1.**
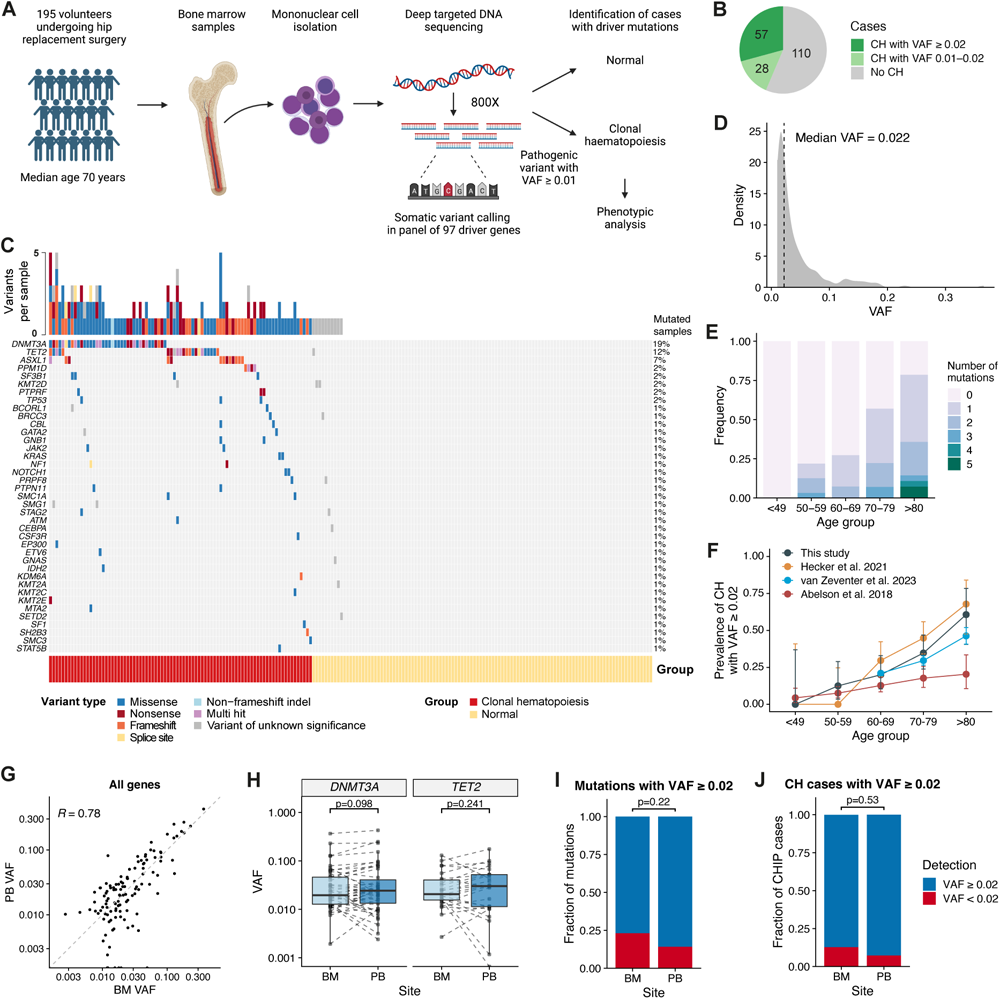
Identification of cases with age-related clonal hematopoiesis in individuals undergoing hip replacement surgery. (A) Experimental design for identifying individuals with clonal hematopoiesis. Bone marrow samples were collected from 195 individuals undergoing hip replacement surgery. Targeted re-sequencing (mean coverage ∼800x) using a 347 kb panel (Table S2) was performed on DNA extracted from BM MNCs to identify somatic driver mutations (Table S2). Cases with clonal hematopoiesis were defined as those with a somatic driver mutation at a VAF of ≥ 0.01. (B) Fraction of samples in the cohort with one or more driver mutations at VAFs of 0.01–0.02 or ≥ 0.02. (C) Landscape of somatic variants observed in the cohort. Each row represents a gene, and each column corresponds to a participant in the study. Top bar plot indicates the number of mutations per sample. Variants are color-coded and classified as pathogenic or variants of unknown significance (VUS) according to criteria specified in Methods. Samples with >1 variant in any one gene are classified as Multiple. Samples with ≥ 1 pathogenic driver mutation were categorized as having clonal hematopoiesis (bottom bar; red). (D) Distribution of VAFs in all mutations observed across the cohort. (E) Frequency of mutations detected per individual by age group. (F) Prevalence of CH with at least one driver mutation (VAF ≥ 0.02) by age. BM DNA sequencing data from participants in this study (n = 195) are compared with a cohort of 676 individuals reported by Abelson et al.^22^, a cohort of 3,359 individuals reported by van Zeventer et al.^58^, and a cohort of 199 individuals undergoing total hip replacement (n = 109 BM samples and n = 91 PB samples) reported by Hecker et al.^59^ Error bars represent 95% confidence intervals. (G) Comparison of VAFs for 128 mutations in paired BM and PB samples. Mutations detected with VAF ≥ 0.01 in either sample type were included. The dashed line shows the line of equality where BM VAF is equal to PB VAF. *R* indicates the Pearson correlation coefficient. (H) Pairwise comparison of VAFs for mutations detected in *DNMT3A* (n = 35) and *TET2* (n = 19). Statistical analysis performed using Wilcoxon signed-rank test. (I) Proportion of mutations detected with a VAF ≥ 0.02 in BM or PB DNA (n = 78 mutations detected with a VAF ≥ 0.02 in either BM or PB in 83 individuals with paired BM and PB sequencing data). (J) Proportion of CH cases with at least one mutation detected with a VAF ≥ 0.02 in BM or PB (n = 83 cases with paired BM and PB sequencing data and a mutation with VAF ≥ 0.02 in either BM or PB).

CH prevalence correlated with increasing age at a VAF cutoff of 0.02 (Figure 1F) or 0.01 (Figure S1C). Comparison with previous studies showed heterogeneity in the frequency of CH cases across studies,^22,58,59^ and the prevalence in our cohort was similar to that observed in a study of peripheral blood (PB) DNA from 3,359 individuals in the general population,^58^ and to another cohort of individuals undergoing hip replacement surgery^59^ (Figure 1F and Figure S1C).^59^ Most studies of CH to date have performed sequencing on peripheral blood (PB) DNA. To determine whether there is any difference in sensitivity for CH detection in BM compared to PB, we compared mutation detection in paired PB granulocytic and BM MNCs DNA on 72 samples with CH and 27 samples without CH. PB sequencing identified an additional 14 mutations not previously called in BM sequencing. Although VAFs differed considerably between BM and PB for some mutations, there was no difference across all mutated genes (Figure 1G; p = 0.11; Wilcoxon signed-rank test), or specifically for *DNMT3A* and *TET2* mutations (Figure 1H). Of 78 mutations detected with a VAF ≥ 0.02 in either BM or PB, 77% were detected in BM with a VAF ≥ 0.02, and 85% were detected in PB with a VAF ≥ 0.02 (Figure 1I). There was also no significant difference in the number of individuals detected with CH between BM and PB (Figure 1J). Our analysis suggests somatic mutations are detected with comparable sensitivity in PB and BM and that the frequency of CH is similar between patients undergoing hip replacement surgery and the general population.

### HSPC differentiation trajectory in DNMT3A and TET2-mutant clonal hematopoiesis

As mutant cells in human CH occur at low frequency, a high-fidelity method for distinguishing mutant and WT cells within the same sample is needed to accurately study the consequences of mutations in human BM. To address this, we further optimized TARGET-seq^60^, which combines high-fidelity single-cell genotyping with transcriptome sequencing on flow cytometry index sorted cells. Our new method, TARGET-seq+, incorporates Smart-seq3 chemistry^61^ to increase transcript detection sensitivity. To validate TARGET-seq+, we compared TARGET-seq and TARGET-seq+ on JURKAT cells and primary human lineage^-^ (Lin^-^) CD34^+^ HSPCs (Figure S1D). Sequencing metrics were comparable between the methods (Figure S1E), but TARGET-seq+ yielded a higher proportion of cells passing quality filters (Figures S1F). Furthermore, TARGET-seq+ increased the number of genes detected per cell by 13.5% in JURKAT cells and by 19.0% in HSPCs (Figures S1G-H). Increased transcript detection sensitivity was observed in both frequently and lowly expressed genes (Figure S1I). Consistent with more efficient transcript capture, cell-to-cell correlations of transcript detection improved significantly with TARGET-seq+ (Figure S1J).

We applied TARGET-seq+ to 9 CH samples and 4 age-matched controls without known driver mutations (non-CH samples) (Figure 2A, Table S3). We focused on hematopoiesis in *DNMT3A* and *TET2*-mutant BM samples by selecting 5 cases with heterozygous *DNMT3A* LoF mutations, 3 with *TET2* LoF mutations, and one case with mutations in both *DNMT3A* and *TET2.* VAFs in BM ranged from 0.061 to 0.366 (Table S3). In two CH cases, there were additional mutations at low VAF (<0.02) (Table S3). We sorted a mean of 1071 cells/sample (range 348–1824), composed of purified Lin^-^CD34^+^ HSPCs, further enriched for primitive Lin^-^ CD34^+^CD38^-^ HSCs and multipotent progenitors (MPPs), combined with CD34^-^ mature cells (Figure S2A). Approximately 40-45% of cells across all samples were Lin^-^CD34^+^CD38^-^; a similar percentage were Lin^-^CD34^+^CD38^+^; and ∼10% were CD34^-^ (Figure S2B). 95.0% of sorted cells (13,247/13,939) were included for further analysis after quality filtering (Figure S2C). We detected ∼10^6^ RNA counts (Figure S2D) and a median of 6484 genes per cell (Figure S2E). These metrics were consistent across samples.

**Figure 2.**
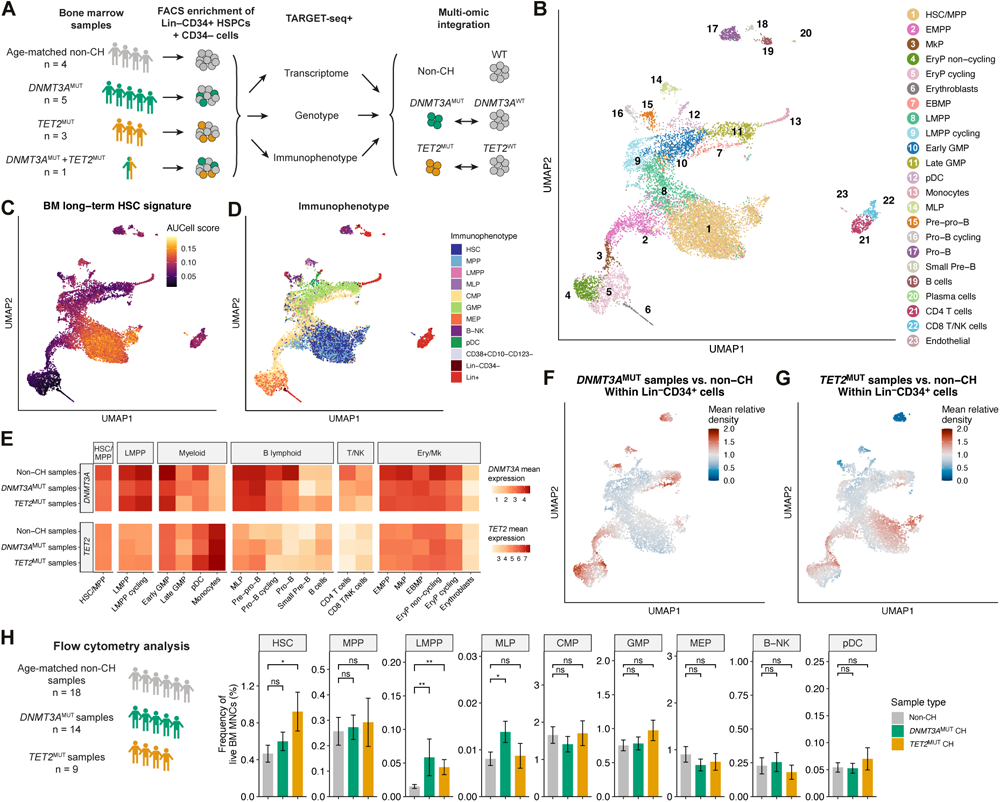
Hematopoietic differentiation trajectory in *DNMT3A* and *TET2*-mutant clonal hematopoiesis. (A) Experimental design for TARGET-seq+ analysis of BM samples from 9 donors with CH and 4 age-matched samples without CH (non-CH). Samples were FACS enriched for Lin^-^ CD34^+^ HSPCs, Lin^-^CD34^+^CD38^-^ cells and CD34^-^ cells and processed using TARGET-seq+ obtaining transcriptome, targeted genotyping, and flow cytometry index immunophenotyping for each cell. (B) UMAP projection of integrated single-cell transcriptome data (n = 13,247 cells from 13 donors). Cells are colored by their cluster annotation, see also Figure S2. (C) UMAP superimposed with AUCell enrichment scores for the BM long-term HSC signature.^70^ (D) UMAP superimposed with the cell immunophenotype determined from flow cytometry indexing. (E) Heatmap of mean log_2_(normalized counts) for *DNMT3A* and *TET2* in control, *DNMT3A*-and *TET2*-mutant samples across hematopoietic cell types. (F and G) UMAPs colored by the mean density of Lin^-^CD34^+^ cells in *DNMT3A*-mutant (F) and *TET2*-mutant (G) CH samples relative to age-matched non-CH samples. Cells sorted from the total Lin^-^CD34^+^ FACS gate were selected (n = 6,629 cells) and MELD ^97^ was used to estimate the density of cells from each sample across the transcriptomic landscape. The mean density for each sample type was then calculated and normalized to the mean density in non-CH samples. A relative density > 1 indicates that the probability of observing a given cell is greater in CH samples than in non-CH samples, whereas a relative density < 1 indicates that the probability is lower in CH samples than in non-CH samples. (H) Flow cytometry analysis was performed on bone marrow samples from normal age-matched non-CH samples (n = 18), and CH samples with either *DNMT3A* (n = 14) or *TET2* (n = 9) mutations present in the largest clone. Barplots show the frequency of stem/progenitor cells as a percentage of BM MNCs. Data are represented as mean ± SEM. P-values calculated by Wilcoxon rank sum test with Holm-Bonferroni multiple testing correction. * p < 0.05, ** p < 0.01. HSC, hematopoietic stem cells; MPP, multipotent progenitor; LMPP, lympho-myeloid primed multipotent progenitor; GMP, granulocyte-monocyte progenitor; pDC, plasmacytoid dendritic cell progenitor; EMPP, erythroid/megakaryocyte primed multipotent progenitor; MkP, megakaryocytic progenitor; EryP, erythroid progenitor; EBMP, Eosinophil/basophil/mast cell progenitor; MLP, multi-lymphoid progenitor; B-NK, B and NK cell progenitor.

To generate a hematopoietic landscape based on single-cell RNA profiles, we integrated gene expression data across the 13 samples (Figure S2F). We annotated 23 cell clusters using published gene signatures^62–70^ and previously described marker genes for specific populations and lineages (Figures 2B, 2C, S2G-H). Using the flow cytometry index data, we overlaid immunophenotypic population identities on the transcriptional landscape (Figure 2D).

Immunophenotypically defined HSPC populations were often present in more than one transcriptionally defined population (Figure S2I), consistent with transcriptional clusters providing a more granular view of hematopoietic cell states, as previously suggested.^62,67^

Downstream of the HSC/MPP cluster, a continuum of transcriptional cell states was observed, with initial separation into two distinct clusters, namely the lymphoid-primed multipotent progenitor (LMPP) and the erythro/megakaryocytic-primed multipotent progenitor (EMPP) (Figure 2B). A series of cell states with neutrophil/monocyte and lymphoid potential extended downstream of LMPPs, whereas progressing from the EMPP were erythroid cells and megakaryocytes, consistent with prior data.^64,71^ By examining *DNMT3A* and *TET2* gene expression across hematopoiesis, we found that both genes were expressed in HSC/MPPs and LMPP, with reduced expression in lymphoid and erythroid lineages (Figure 2E). Notably, *TET2* expression increased during myeloid maturation, in contrast to *DNMT3A*.

Even though CH is mainly composed of normal cells, prior data have shown the frequencies of immunophenotypic HSPC change with age^72^ and in myeloid disease.^73–76^ As a foundation to examine hematopoiesis in CH, we used two approaches to document the size of the HSPC compartments in CH. First, we measured the frequency of cells across the transcriptional landscape and compared CH samples with non-CH samples (details in Methods). On average, *DNMT3A*-mutant samples had a modest 0.89-fold reduction in transcriptional HSC/MPPs and early lympho-myeloid progenitors, and a 1.2 to 1.4-fold increase in late progenitor cells (Figure 2F). By contrast, *TET2*-mutant samples showed a modest 1.2-fold expansion of transcriptional HSC/MPPs (Figure 2G). We then performed conventional immunophenotyping on a larger sample set, which showed Lin^-^CD34^+^CD38^-^CD45RA^-^CD90^+^ HSCs and CD49f^+^ long-term HSCs (LT-HSC) were 2-fold and 2.2-fold expanded, respectively, in *TET2*-mutant samples relative to non-CH samples (Figures 2H and S2J). Additionally, *DNMT3A-* and *TET2*-mutant CH samples showed a 3.9-fold and 2.9-fold increase in the rare immunophenotypic Lin^-^ CD34^+^CD38^-^CD45RA^+^ LMPP population respectively (Figure 2H), which constitutes a minor fraction of the transcriptionally-defined LMPP and multi-lymphoid progenitors (MLP) (Figure S2I). Overall, the sizes of the HSPC compartments are moderately perturbed in *DNMT3A*-and *TET2*-mutant CH.

### Distinct patterns of clonal expansion of DNMT3A-and TET2-mutant clones

Global differences in HSPC compartments fail to reveal the differential size of co-existing CH mutant clones as opposed to WT cells. To address this, we directly compared mutant and WT cells within the same samples, by integrating single-cell genotyping data with gene expression profiles. *DNMT3A* and *TET2* loci were successfully amplified in 97.7% and 98.1% of cells respectively, resulting in clonal assignment for 93.1% of CH cells (Figure S3A). Importantly, allelic dropout (ADO) rates were low (7.2–13.4% across 4 loci in which ADO was estimated by a heterozygous germline single nucleotide polymorphism in the genotyping amplicon), allowing us to confidently identify most mutant and WT cells (Figure S3B). Genotyping rates for *DNMT3A* and *TET2* were high across all cell types (Figure S3C), and the frequency of mutant cells detected by TARGET-seq+ correlated well with estimates from bulk BM DNA sequencing (Figure S3D). In all the subjects with more than one clone, *DNMT3A* or *TET2* mutations were present in the founder clone, allowing us to determine the effect of these mutations in a WT background (Figures S3E-L).

To determine the expansion or contraction of mutant clones through hematopoietic differentiation, we projected genotypes onto the transcriptomic differentiation landscape and compared the density of mutant and wild type cells (Figure 3A). We included all successfully genotyped cells from CH samples (including Lin^-^CD34^+^, Lin^-^CD34^+^CD38^-^ and CD34^-^ sorting strategies). By considering the ratio of cell density between mutant and WT cells (hereafter referred to as the mutant clone likelihood) and normalizing this likelihood to that of the HSC/MPP cluster within each sample, we could determine how clone size changes downstream of the HSC/MPP (Figure 3A).

**Figure 3.**
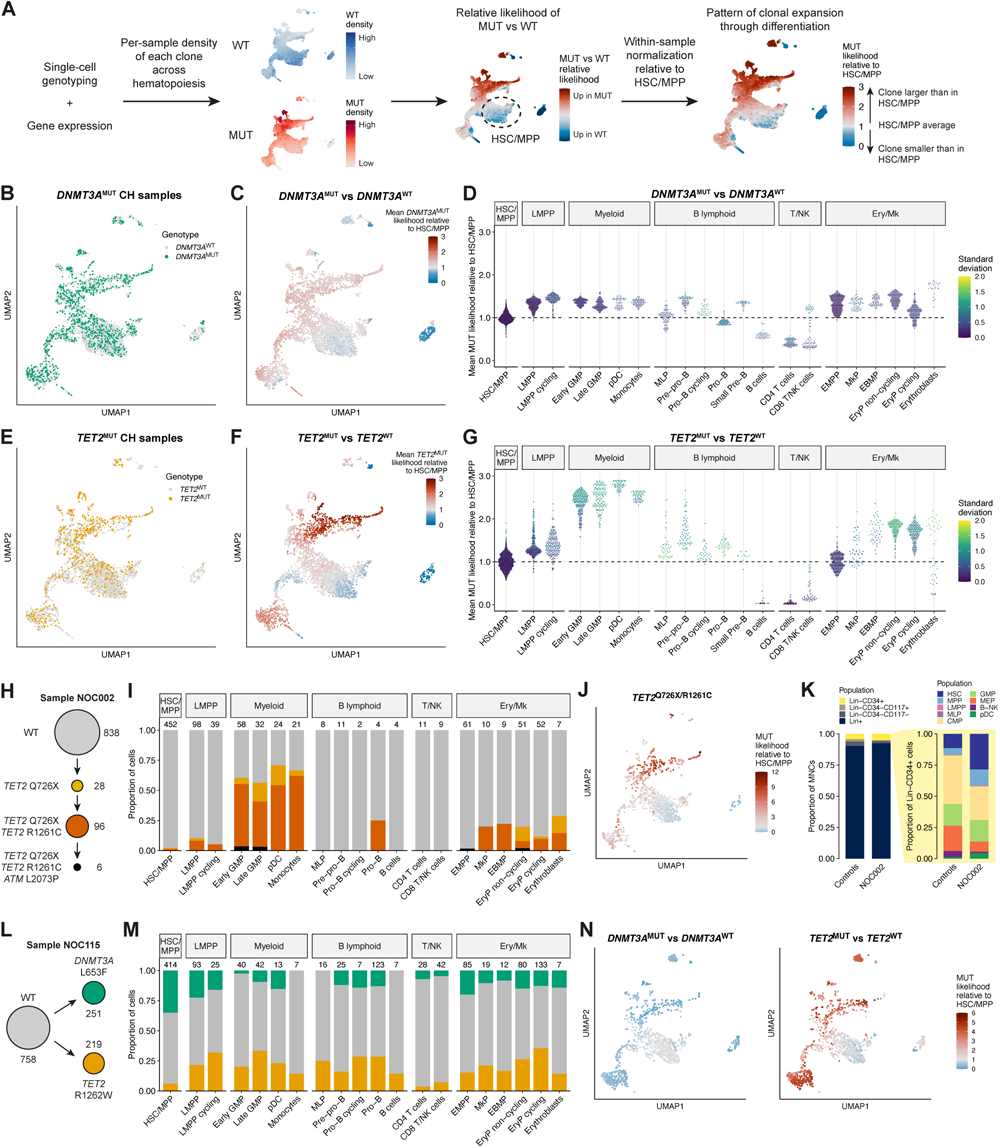
Distinct patterns of clonal expansion of *DNMT3A-* and *TET2*-mutant clones. (A) Single-cell genotyping and gene expression data were integrated and, for each sample, MELD was used to estimate the density of cells from each clone across the transcriptomic landscape (see Methods). The relative likelihood of a cell being mutant vs. WT (relative clone likelihood) was calculated and then normalized to the mean likelihood in the HSC/MPP cluster within each sample. A mutant relative likelihood > 1 indicates that the probability of a given cell being mutant is greater than in the HSC/MPP, whereas a relative likelihood < 1 indicates that the probability is lower than in the HSC/MPP. (B) UMAP of WT and single-mutant cells from *DNMT3A*^MUT^ CH samples (n = 4301 cells from 5 samples) colored by genotype, either wild-type (WT, n = 2813) or *DNMT3A*^MUT^ (n = 1488). (C) UMAP of WT and single-mutant cells from *DNMT3A*^MUT^ CH samples colored by the mean likelihood of cells being *DNMT3A*^MUT^ relative to the average within HSC/MPP. The mean value across 5 samples is shown. (D) Scatterplot showing the distribution of mean *DNMT3A*^MUT^ clone likelihoods by cluster. Each dot represents a cell. The y axis shows the mean *DNMT3A*^MUT^ clone likelihood relative to the HSC/MPP as in (C). Each dot is colored by the standard deviation of the values across 5 samples. (E) UMAP of WT and single-mutant cells from *TET2*^MUT^ CH samples (n = 3472 cells from 3 samples) colored by genotype, either wild-type (WT, n = 2655) or *TET2*^MUT^ (n = 817). (F) UMAP of WT and single-mutant cells from *TET2*^MUT^ CH samples colored by the mean likelihood of cells being *TET2*^MUT^ relative to the average within HSC/MPP. The mean value across 3 samples is shown. (G) Scatterplot showing the distribution of mean *TET2*^MUT^ clone likelihoods by cluster. Each dot is colored by the standard deviation of the values across the 3 samples. (H) Clonal structure for the NOC002 sample in which 2 separate *TET2* mutations and an *ATM* mutation were detected. Cell numbers assigned to each clone are indicated. (I) Clonal composition within each cluster for sample NOC002. Each clone is colored as in (H). The number of cells analyzed in each cluster is shown above. (J) UMAP showing the likelihood of cells being in the double *TET2*^MUT^ clone (*TET2*^Q726X/R1261C^) relative to the average within HSC/MPP in the NOC002 sample. (K) Immunophenotypic BM compartment sizes for the NOC002 sample. Left-hand bars show compartments as a proportion of total BM MNCs, while right-hand bars show HSPC compartments within Lin^-^CD34^+^ cells. Data from the NOC002 sample are compared with the median population sizes from 18 age-matched control samples. (L) Clonal structure for the NOC115 sample in which *DNMT3A* and *TET2* mutations were detected. The two mutations were mutually exclusive in single-cell genotyping. Cell numbers assigned to each clone are indicated. (M) As in (I) but for sample NOC115. Each clone is colored as in the clonal structure in (L). (N) UMAPs showing the likelihood of cells being in the *DNMT3A*^MUT^ (left) and *TET2*^MUT^ (right) clones relative to the average within HSC/MPP in the NOC115 sample.

First, we analyzed the pattern of expansion of clones with a single *DNMT3A* mutation. Cells with a single *DNMT3A* mutation were intermingled with *DNMT3A*^WT^ cells, both within individual samples (Figures S3E-I) and in the integrated dataset (Figure 3B), indicating that *DNMT3A*^MUT^ and *DNMT3A*^WT^ cells shared a similar differentiation trajectory and that mutant cells did not create novel cell states. In HSC/MPPs, the contribution of mutant cells was highly variable, with the clone size ranging from 3.4% to 70.4% (Figures 3B and S3E-I). Across all samples, changes in *DNMT3A*^MUT^ clone size through differentiation were modest (Figures 3C and 3D). Downstream of HSC/MPPs, the mean mutant clone likelihood was approximately 50% higher in early EMPP and LMPP populations than in HSC/MPPs (Figures 3C and 3D). Clone sizes were then largely maintained at later stages of differentiation, except for in T cells and to a lesser extent B cells, where clone size was on average 50% smaller than in HSC/MPPs. Depletion of *DNMT3A*^MUT^ cells in lymphoid cells was variable between individuals, with mutant cells being nearly absent from lymphoid cells in one sample (Figure S3E), consistent with previous data from purified peripheral blood cell populations.^77–79^ Overall, *DNMT3A*^MUT^ clonal expansion occurred primarily in HSCs and early multipotent progenitors. Furthermore, aside from depletion in B and T cells, there was no evidence of lineage bias in *DNMT3A*^MUT^ cells, in contrast to data from myeloma remission samples in which *DNMT3A*^R882^–mutant CH cells were biased towards the megakaryocytic-erythroid lineage.^56^

*TET2*^MUT^ cells also intermingled with *TET2*^WT^ cells (Figure 3E). Clone size within HSC/MPPs varied between 1.1%–32.6% (Figures 3E and S3J-L). In contrast to *DNMT3A*^MUT^ clones, there was more pronounced expansion of *TET2*^MUT^ clones downstream of HSC/MPPs during myelopoiesis (Figures 3F and 3G). Indeed, *TET2*^MUT^ clones were on average 2.5–3 fold larger in granulocyte-monocyte progenitors (GMP) compared to HSC/MPPs. In 2 out of 3 samples, the mutant clone also expanded within erythroid progenitors (Figures S3K and S3L). By contrast, *TET2*^MUT^ cells were almost absent from mature B and T cells in all samples, suggesting inability to complete terminal lymphoid differentiation. Conversely, there was heterogeneity in the contribution of *TET2*^MUT^ cells to earlier lymphoid progenitors. In 2/3 cases we observed a depletion of mutant cells from these populations (Figures S3J and S3L), but in one individual the *TET2*^MUT^ clone constituted 91% of B cell progenitors (Figure S3K).

Next, to understand the differentiation potential of heterozygous versus homozygous *TET2*^MUT^ clones, we studied one sample where two *TET2* mutations were acquired sequentially in a linear clonal structure, followed by acquisition of an *ATM* mutation in a small terminal subclone (Figure 3H). The single-and double-*TET2*^MUT^ clones each contributed to only 1.1% of HSC/MPPs. While the single-*TET2*^MUT^ clone expanded 3–4 fold in downstream erythroid and myeloid progenitors, the double-*TET2*^MUT^ clone dramatically outcompeted the single mutant clone during myelopoiesis, contributing to more than half of all cells in GMP, plasmacytoid dendritic cells (pDC) and monocytes (Figures 3I and 3J). Notably, immunophenotypic pDC were expanded in this sample (Figure 3K). Interestingly, there was also a 2.6-fold expansion of the immunophenotypic HSC and MPP populations in this sample relative to non-CH samples (Figure 3K). Since 97.6% of transcriptional HSC/MPPs were *TET2*^WT^, this raises the question of whether *TET2*^MUT^ CH might increase WT HSC/MPP cell numbers in a non-cell autonomous manner (Figure 3K). Overall, *TET2* LoF leads to a competitive advantage within HSCs and biased cell output towards myelopoiesis in a gene dose dependent manner, consistent with *TET2* gene expression across hematopoiesis (Figure 2E).

Finally, we had the informative opportunity to study clonal competition between *DNMT3A*^MUT^ and *TET2*^MUT^ cells in the same individual in a sample with co-existing, independently acquired *DNMT3A*^MUT^ and *TET2*^MUT^ clones (Figure 3L). Interestingly, the *DNMT3A*^MUT^ clone was 6 times larger than the *TET2*^MUT^ clone within HSC/MPPs, but the *TET2*^MUT^ clone became 4 times larger than the *DNMT3A*^MUT^ clone within the GMPs (Figure 3M and 3N). Notably, the *TET2*^MUT^ clone was also larger than the *DNMT3A*^MUT^ clone in erythroid and early lymphoid progenitors, as well as during early B cell lymphopoiesis (Figure 3M).

In summary, *DNMT3A*^MUT^ and *TET2*^MUT^ clones showed distinct patterns of clonal expansion across differentiation. The selective advantage of *DNMT3A*^MUT^ clones occurs mainly in HSCs and early multipotent progenitors, whereas *TET2*^MUT^ clones expand not only in HSCs, but also further through differentiation, especially in myelopoiesis.

### Transcriptional basis for dysregulated myeloid differentiation of TET2-mutant clones

As *TET2*^MUT^ clonal expansion was most pronounced in the myeloid lineage, we further explored the transcriptional basis of this phenotype in individuals with *TET2*^MUT^ CH. We first compared the density distributions of *TET2*^MUT^ and *TET2*^WT^ cells along myeloid pseudotime (the differentiation trajectory from HSCs to mature myeloid cells) (Figure 4A). *TET2*^MUT^ cells accumulated particularly at the progenitor stage, within the cycling LMPP, early GMP, and late GMP transcriptional clusters, and to a lesser extent at later stages of the trajectory, in mature plasmacytoid dendritic cell (pDC) and monocyte clusters (Figure 4B).

**Figure 4.**
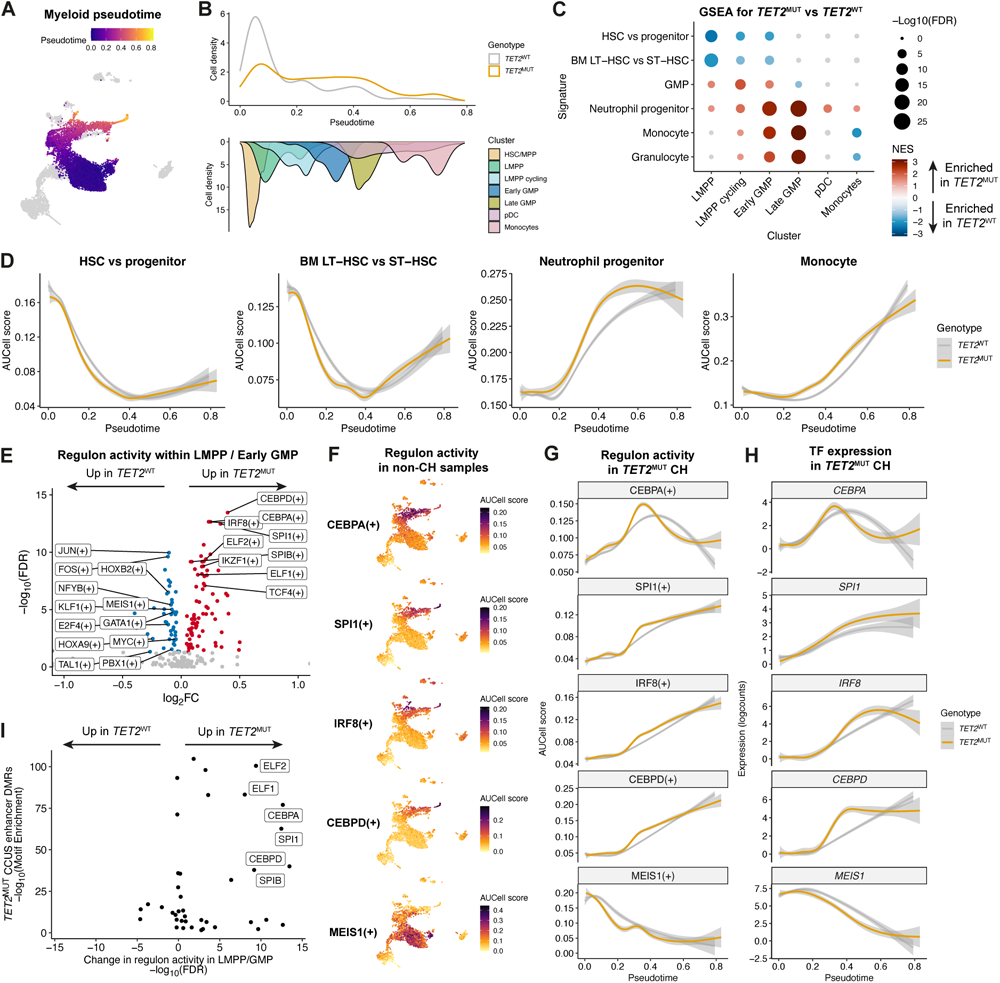
*TET2*-mutant clones lead to dysregulated myeloid differentiation. (A) UMAP showing the myeloid differentiation trajectory with cells colored by pseudotime score. (B) Top: Density plot showing the distribution of *TET2*^WT^ and *TET2*^MUT^ cells through pseudotime in the myeloid lineage. Cells sorted as part of the total Lin^-^CD34^+^ FACS gate were downsampled to an equal number cells per sample (n = 176 cells from each of 4 samples). Bottom: Histogram showing the density of cells in each cluster along pseudotime for cells included in this analysis. (C) GSEA against hematopoietic signatures comparing *TET2*^MUT^ versus *TET2*^WT^ cells within each LMPP and GMP cluster. Differential expression analysis was performed accounting for sample and batch effects. Cells from the 4 *TET2*^MUT^ CH samples were included in the analysis. Color intensity indicates the normalized enrichment score (NES); signatures with FDR > 0.05 are colored grey. Positive NES values indicate enrichment in mutant cells. BM, bone marrow; LT-HSC, long-term HSC; ST-HSC, short-term HSC. (D) Local regression of AUCell expression scores for HSC and myeloid gene signatures along myeloid pseudotime, comparing *TET2*^WT^ and *TET2*^MUT^ cells. (E) Volcano plot showing differentially expressed regulons between *TET2*^MUT^ and *TET2*^WT^ cells within the LMPP cycling and early GMP clusters in *TET2*^MUT^ CH samples. FDR-corrected P-values calculated by linear mixed model test accounting for sample effects. (F) UMAPs showing activity of the indicated regulons across the hematopoietic landscape within non-CH samples. (G) Local regression of regulon activity through myeloid pseudotime, comparing *TET2*^MUT^ and *TET2*^WT^ cells in *TET2*^MUT^ CH samples. (H) Fitted gene expression values along pseudotime for the transcription factors shown in (F) and (G) in *TET2*^MUT^ and *TET2*^WT^ cells in the myeloid lineage. I) Enrichment of TF motifs within differentially methylated enhancer regions (DMRs) that are hypermethylated in monocytes from *TET2*-mutant CCUS patients^81^ (y-axis), plotted against the ranked change in regulon activity (–log10(FDR) * sign of the fold change) between *TET2*^MUT^ and *TET2*^WT^ cells within the LMPP cycling and early GMP clusters from (E).

At least two reasons may explain the increased ratio of *TET2*^MUT^ versus *TET2*^WT^ cells within progenitor stages of myelopoiesis. *TET2*^MUT^ myeloid progenitor expansion might arise either from reduced retention of stem cell transcriptional programs and increased myeloid differentiation from HSCs, and/or due to altered maturation of myeloid progenitors.

To begin to address this, we performed gene set enrichment analysis (GSEA) using published HSPC signatures^62,66,70^ to examine transcriptional differences between *TET2*^MUT^ and *TET2*^WT^ cells within LMPP and GMP clusters. We first validated these signatures in our dataset (Figure S4A). Compared to *TET2*^WT^ cells, *TET2*^MUT^ LMPPs, cycling LMPPs and GMPs were negatively enriched for HSC signatures (Figure 4C). Conversely, *TET2*^MUT^ GMPs were enriched for neutrophil progenitor and mature neutrophil/monocyte signatures (Figure 4C). Concordantly, CD38 and CD45RA surface protein expression was higher in *TET2*^MUT^ compared to *TET2*^WT^ LMPPs (Figure S4B), consistent with our previous data that higher expression of these markers enriches for myeloid potential in LMPPs.^67^ Finally, megakaryocytic-erythroid signatures were also negatively enriched, particularly in *TET2*^MUT^ cycling LMPP cells (Figure S4C), consistent with their myeloid bias.

To further explore differentiation kinetics, we analyzed the same HSPC signatures across myeloid pseudotime by computing the AUCell score^80^ for these signatures in *TET2*^MUT^ and *TET2*^WT^ cells (Figure 4D). This showed that HSC genes (both genes with increased expression in HSCs versus CD34^+^ progenitors and those with increased expression in BM LT-HSC versus short-term HSCs (ST-HSC)) showed a faster reduction in expression in *TET2*^MUT^ compared to *TET2*^WT^ cells along the trajectory. Furthermore, gene sets associated with neutrophil progenitors and monocytes showed earlier, or “premature”, expression in *TET2*^MUT^ cells. Concordantly, exemplar genes expressed in mature myeloid cells, including *MPO*, *NKG7*, *KLF4*, and *RBM47*, showed premature expression in *TET2*^MUT^ progenitors along myeloid pseudotime (Figure S4D). Taken together, this suggests that early *TET2*^MUT^ lympho-myeloid progenitors retain less HSC and non-myeloid programs, while later in differentiation, *TET2*^MUT^ progenitors commit more rapidly to myelopoiesis.

We next wanted to identify potential drivers of earlier expression of mature myeloid gene programs in *TET2*^MUT^ myeloid progenitors. We used pySCENIC to compare expression of TFs and their putative downstream targets genes (i.e. regulons) between *TET2*^MUT^ and *TET2*^WT^ cells within LMPP and early GMP clusters, where lymphoid and myeloid lineages diverge. The canonical myeloid TFs CEBPD, CEBPA, IRF8 SPI1, and SPIB were all more active in *TET2*^MUT^ LMPPs and early GMPs (Figure 4E). Conversely, TFs associated with HSC self-renewal (MEIS1, HOXA9, HOXB2, NFYB and PBX1), and with megakaryocytic-erythroid differentiation (GATA1, TAL1, and KLF1) were less active in *TET2*^MUT^ cells. Regulon activity (Figure 4F and 4G), as well as TF expression (Figure 4H) of *CEBPA*, *SPI1*, *IRF8*, and *CEBPD*, peaked earlier in *TET2*^MUT^ versus *TET2*^WT^ cells along myeloid pseudotime. This quartet of myeloid TFs are required both in early (CEBPA, SPI1 and IRF8) and later (SPI1, IRF8 and CEBPD) stages of myelopoiesis. In contrast, expression of *MEIS1* and its targets were downregulated earlier in *TET2*^MUT^ cells. Interestingly, binding motifs of CEBPA, CEBPD, SPI1, SPIB, and ELF1/2 were also enriched within differentially methylated enhancers in peripheral blood granulocytes from patients with *TET2*^MUT^ cytopenia of undetermined significance (CCUS) (Figure 4I).^81^ This suggests a link between altered enhancer methylation and dysregulated myeloid TF activity in *TET2*^MUT^ myeloid cells.

Finally, we explored the transcriptional consequences of earlier and aberrant myeloid differentiation in *TET2*^MUT^ cells. When we compared *TET2*^MUT^ and *TET2*^WT^ LMPP and GMP, and performed GSEA (Figure S4E), we identified enrichment of signatures associated with cell cycle (cycling LMPP), oxidative phosphorylation, cytokine signaling, and innate immune effector function (GMP) in *TET2*^MUT^ cells. Overall, this suggests that *TET2*^MUT^ myeloid progenitors are biased towards maturation, with accelerated upregulation of mature gene expression programs and dysregulated expression of inflammatory pathways.

### Non-cell-autonomous activation of inflammatory transcriptional programs in HSC/MPP in clonal hematopoiesis is attenuated in mutant HSCs

Humans have an estimated 50,000–200,000 HSCs.^82^ In an individual in which 1% of HSCs harbor a CH mutation, this would represent a 500–2,000-fold expansion from a single initiating mutant HSC. When compared to this, changes in clone size observed downstream of the HSC compartment are modest, implying that the greatest clonal expansion for both *DNMT3A*^MUT^ and *TET2*^MUT^ (CH^MUT^) clones occurred within long-lived HSC/MPPs. At least two hypotheses could explain the relative clonal advantage of CH^MUT^ HSCs over WT (CH^WT^) HSCs: either CH^MUT^ HSCs may have a cell-autonomous competitive advantage and/or CH^WT^ HSCs may be at a competitive disadvantage.

To begin to dissect which hypotheses were operative, we first compared gene expression between HSC/MPPs from CH samples (both CH^MUT^ and CH^WT^) and HSC/MPPs from age-matched samples without CH (non-CH samples) (Figure 5A). We first identified differentially expressed genes between either *DNMT3A*^MUT^ (Figure 5B top left) or *TET2*^MUT^ (Figure 5B top right) HSC/MPPs and non-CH HSC/MPPs. GSEA showed enrichment of TNFα signaling via NF-κB, inflammatory response, and HSC quiescence signatures in both *DNMT3A*^MUT^ and *TET2*^MUT^ HSC/MPPs compared to non-CH HSC/MPPs. By contrast, gene sets for progenitors (GMP and to a lesser extent LMPP) were negatively enriched in *DNMT3A*^MUT^ and *TET2*^MUT^ HSC/MPPs, suggesting CH^MUT^ HSC/MPPs are less primed towards differentiation. Strikingly, a similar pattern of enrichment was observed when *DNMT3A*^WT^ and *TET2*^WT^ HSC/MPPs from CH samples were compared to non-CH HSC/MPPs (Figure 5B, bottom panels). Taken together, these transcriptional data suggest HSC/MPPs in CH individuals are impacted by an inflammatory milieu, regardless of whether they have *DNMT3A* or *TET2* mutations.

**Figure 5.**
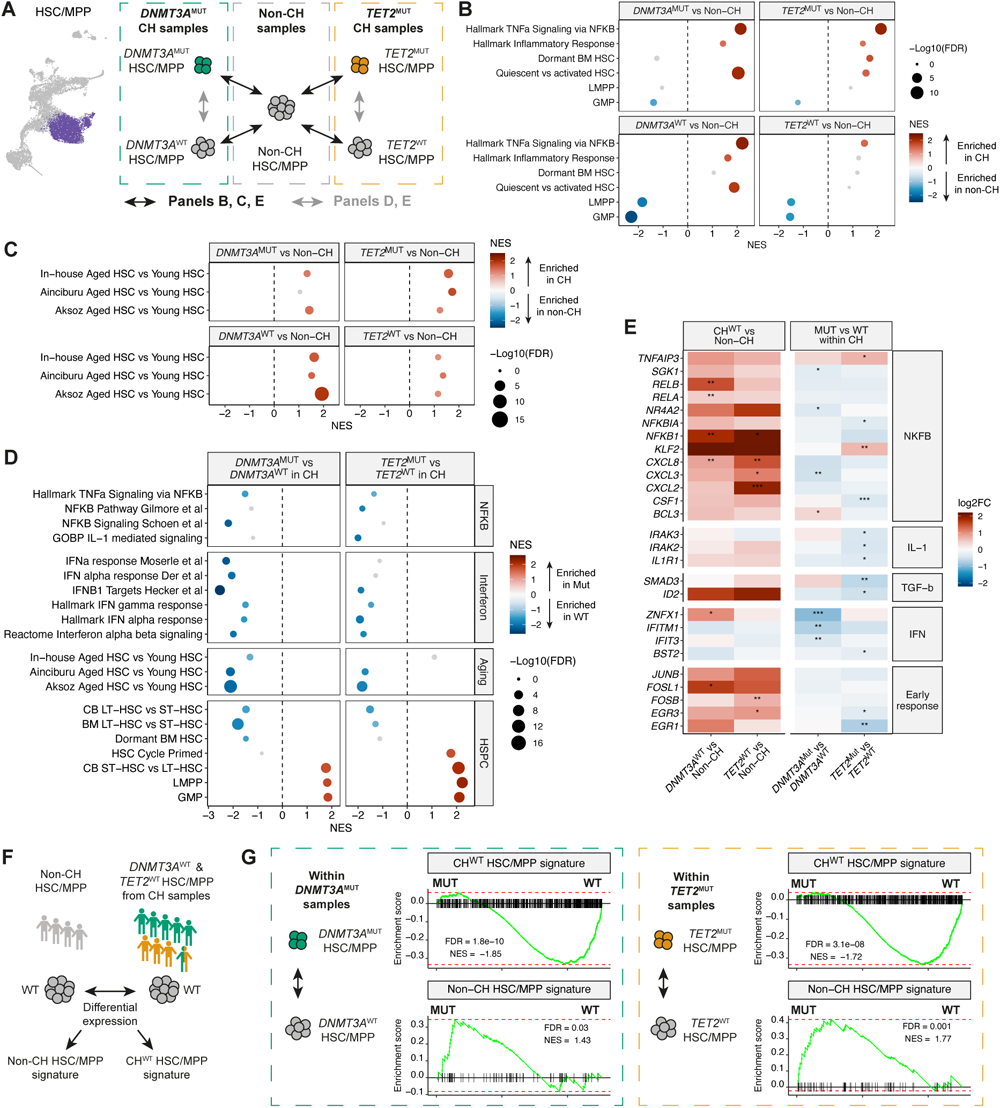
Non-cell-autonomous activation of inflammatory transcriptional programs in HSC/MPP in clonal hematopoiesis is attenuated in mutant HSCs. (A) Schematic showing the strategy for differential gene expression analysis between HSC/MPPs from CH samples and HSC/MPPs from age-matched non-CH samples (black arrows; panels B, C, E), and between CH^MUT^ and CH^WT^ HSC/MPPs within CH samples (grey arrows; panels D, E). (B) GSEA against Hallmark inflammatory signatures and hematopoietic lineage signatures comparing WT or mutant HSC/MPPs from CH samples with HSC/MPPs from age-matched non-CH samples. Differential expression analysis was performed accounting for sample, age, and batch effects. The left panels show comparisons between the 4 non-CH samples (n = 1280 cells) and the five *DNMT3A*^MUT^ samples (n = 1145 WT cells, n = 405 *DNMT3A*^MUT^ cells). The right panels show comparisons between the 4 non-CH samples (n = 1280 cells) and the three *TET2*^MUT^ samples (n = 1243 WT cells, n = 221 *TET2*^MUT^ cells). Note the double-mutant NOC115 sample was excluded from this analysis. Signatures with FDR > 0.2 are colored grey. Positive NES values indicate enrichment in cells from CH samples. (C) As in (B) but showing GSEA against aged HSC signatures derived from the in-house dataset, and two additional studies comparing aged and young human HSCs.^86,87^ (D) GSEA against NF-κB, interferon, and hematopoietic signatures comparing *DNMT3A*^MUT^ versus *DNMT3A*^WT^ HSC/MPPs (left) and *TET2*^MUT^ versus *TET2*^WT^ HSC/MPPs (right) within CH samples. Color intensity indicates the normalized enrichment score (NES); signatures with FDR > 0.2 are colored grey. Positive NES values indicate enrichment in mutant cells. BM, bone marrow; CB, cord blood. (E) Heatmap showing log2 fold change in expression of genes related to inflammatory pathways within HSC/MPP. The left panel shows results when comparing WT cells from CH samples vs. cells from non-CH samples. The right panel shows results when comparing mutant cells from CH samples vs WT cells from CH samples. Asterisks represent FDR-corrected p-values from differential expression testing. * FDR < 0.1, ** FDR < 0.05, *** FDR < 0.01. (F) Schematic showing the strategy for deriving CH^WT^ HSC/MPP and non-CH HSC/MPP signatures. Differential expression analysis was performed between HSC/MPPs from the 4 non-CH samples (n = 1280 cells) and WT cells from the 9 CH samples (n = 2632 cells), accounting for sample, age, and batch effects. Genes with FDR < 0.1 and log2FC > 0.5 were included in each signature. (G) GSEA enrichment plots for the CH HSC/MPP signature (top panels) and non-CH HSC/MPP signature (bottom panels), comparing WT and mutant cells within CH samples. Positive enrichment scores indicate enrichment in mutant cells.

Prior data indicate that aging is associated with chronic inflammation, increased NF-κB signaling, increased quiescence and functional decline in HSCs.^83–85^ To evaluate if aging-related signatures were enriched in CH HSC/MPPs compared to age-matched non-CH HSC/MPPs, we performed single-nucleus RNA-seq (snRNA-seq) on human bone marrow HSPCs collected from young and older aged individuals and performed differential expression between HSCs to define signatures of aged HSCs (Figure S5A-D). For further validation, we defined additional signatures of aged HSCs through re-analysis of two additional human HSPC scRNA-seq datasets (Figures S5E-H and Methods).^86,87^ Notably, transcriptional signatures specific to aged human HSCs were enriched in *DNMT3A*^MUT^ and *TET2*^MUT^ HSC/MPPs from CH samples compared to non-CH WT HSC/MPPs, as well as in *DNMT3A*^WT^ and *TET2*^WT^ HSC/MPPs from CH samples compared to non-CH WT HSC/MPPs (Figure 5C).

We then asked how *DNMT3A* and *TET2* mutations alter gene expression to provide CH^MUT^ HSC/MPPs a fitness advantage. Specifically, we wanted to understand the cell-intrinsic effects of the CH mutation. We compared gene expression between CH^MUT^ and CH^WT^ cells and used GSEA to identify gene expression programs that were differentially enriched between *DNMT3A*^MUT^ and *DNMT3A*^WT^ HSC/MPPs (Figure 5D, left) or between *TET2*^MUT^ and *TET2*^WT^ HSC/MPPs (Figure 5D, right). TNFα signaling, NF-κB pathway, IL-1 signaling, and IFNα response signatures were all negatively enriched in CH^MUT^ HSC/MPPs compared to CH^WT^ HSC/MPPs. Furthermore, aged HSC signatures and those that distinguish LT-HSC from ST-HSC were also negatively enriched in CH^MUT^ HSC/MPPs compared to CH^WT^ HSC/MPPs, particularly in individuals with *DNMT3A*^MUT^ CH. Specific genes associated with interferon, NF-κB, IL-1, TGF-ý signaling and early response were more highly expressed in *DNMT3A*^WT^ and *TET2*^WT^ HSC/MPPs from CH individuals compared to HSC/MPPs from non-CH individuals (Figure 5E, left panel). Conversely, many of the same genes were expressed at lower levels in *DNMT3A*^MUT^ and *TET2*^MUT^ HSC/MPPs compared to their CH^WT^ counterparts (Figure 5E, right panel), consistent with CH^MUT^ HSC/MPPs having an attenuated response to the inflammatory environment.

In contrast, signatures of cycle primed HSC, LMPP, and GMP were enriched in CH^MUT^ HSC/MPPs compared to CH^WT^ HSC/MPPs (Figure 5D). Consistent with reduced quiescence (or a more ST-HSC-like phenotype), *DNMT3A*^MUT^ and *TET2*^MUT^ HSC/MPPs also showed positive enrichment for pathways related to mitosis, cell migration, and signaling, particularly in *TET2*^MUT^ cells, compared to CH^WT^ HSC/MPPs (Figure S5I). Furthermore, *TET2*^MUT^ HSC/MPPs had greater RNA content, were larger, more granular, and had lower CD49f protein expression compared to *TET2*^WT^ cells (Figures S5J-M). These results suggest an inverse relationship between molecular programs underlying inflammation and aging in contrast to programs underlying lympho-myeloid differentiation priming within the HSC/MPP compartment. We assessed the expression of these signatures in HSC/MPPs at the single-cell level using AUCell scores. Indeed, expression of TNFα via NF-κB signaling was strongly correlated with the aged HSC signatures, while both sets of signatures were negatively correlated with the LMPP signature (Figure S5N).

These results suggested that CH^MUT^ HSC/MPPs have an altered transcriptional response to the CH environment. To generate molecular signatures that capture those processes that differ between CH and non-CH samples, we compared gene expression profiles of CH^WT^ HSC/MPPs to non-CH HSC/MPPs (Figures 5F and S5O). There were 561 genes upregulated in CH^WT^ HSC/MPPs, whereas only 61 genes were upregulated in non-CH HSC/MPPs (Figure S5O). Genes upregulated in CH^WT^ HSC/MPPs were enriched for TNFα via NF-κB and TGF-β signaling, as well as signatures of HSC quiescence and aging (Figure S5P). We then asked whether *DNMT3A*^MUT^ and *TET2*^MUT^ HSC/MPPs were impacted differently than CH^WT^ HSC/MPPs. Indeed, both *DNMT3A*^MUT^ and *TET2*^MUT^ HSC/MPPs were negatively enriched for the CH^WT^ HSC/MPP signature (Figure 5G top panels), but positively enriched for the non-CH HSC/MPP signature (Figure 5G bottom panels), consistent with our findings above.

In summary, our data suggest that, compared to their CH^WT^ counterparts, CH^MUT^ HSC/MPPs are shifted towards a state that is more similar to non-CH HSC/MPPs. Collectively, these data suggests that the transcriptional response to aging and inflammation may be attenuated in CH^MUT^ HSC/MPPs.

### HSC are transcriptionally heterogeneous where distinct HSC subsets show differing responses to the CH environment

Transcriptional differences between HSC/MPPs in a CH and non-CH context point to a need for a more granular exploration of HSC/MPP heterogeneity. Thus, we subclustered HSC/MPPs and the earliest progenitors from CH and non-CH samples (Figure 6A and Figure S6A), using self-assembling manifolds (SAM), an unsupervised approach to prioritize biologically relevant features among comparatively homogenous cells which has previously been applied to human HSC/MPPs.^88–90^ This identified three distinct HSC clusters (HSC 1-3), separated from the MPP, LMPP, and EMPP clusters. HSC1 and HSC2 were composed of similar numbers of cells (1701 and 2150 respectively); HSC3 had far fewer cells (484). The HSC clusters were the most highly enriched for HSC signatures (including LT-HSC) and depleted for ST-HSC, LMPP, GMP and MEP signatures (Figure 6B and Figure S6B). HSC1 and HSC2 clusters were more enriched for CD49f+ HSCs compared to HSC3 cluster, which was immunophenotypically more similar to MPPs (Figure S6C).

**Figure 6.**
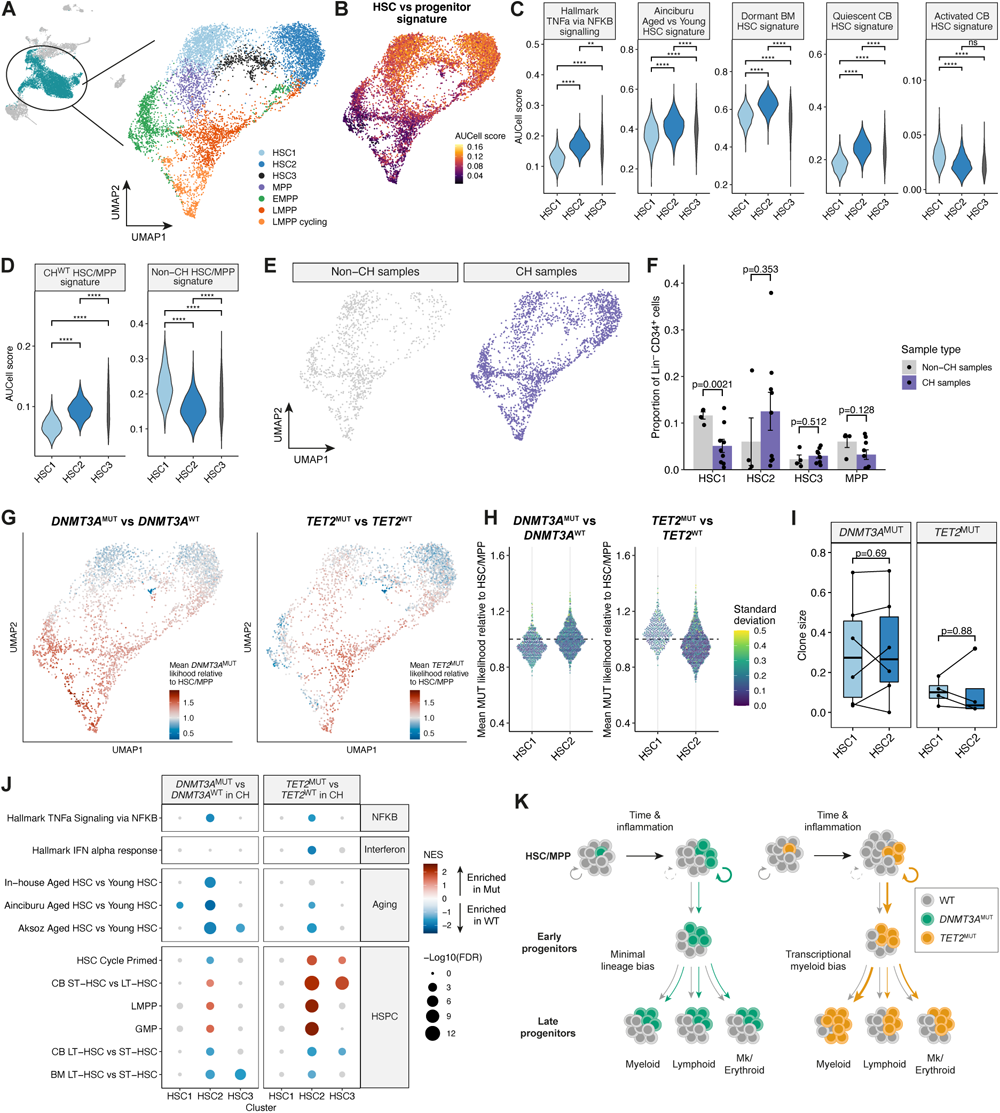
HSC are transcriptionally heterogeneous where distinct HSC subsets show differing responses to the CH environment. (A) UMAP embedding of 8059 cells from the HSC/MPP, EMPP, LMPP and LMPP cycling clusters after feature weight derivation with the Self-Assembling Manifolds (SAM) algorithm. Cells are colored by cluster annotation. (B) UMAP superimposed with AUCell enrichment scores for a signature of genes differentially expressed between HSCs and progenitors^66^. (C) AUCell enrichment scores for the Hallmark TNFa via NF-κB, HSC aging, dormant BM HSC^70^, and quiescent vs activated CB HSC signatures^90^, comparing the 3 HSC clusters. P-values calculated by pairwise unpaired *t* test. The area of each violin is proportional to the cell numbers. (D) AUCell enrichment scores for the CH^WT^ and non-CH HSC/MPP signatures, comparing the 3 HSC clusters. P-values calculated by pairwise unpaired *t* test. (E) UMAP embeddings showing cells from non-CH and CH samples. Cells are colored by sample type. (F) Quantification of the size of each HSC/MPP cluster as a proportion of Lin^-^CD34^+^ cells, comparing CH and non-CH samples. Only cells sorted from the total Lin^-^CD34^+^ FACS gate were included. Data are represented as mean ± SEM. Each dot represents a sample. P-values calculated by unpaired *t* test. (G) UMAPs of cells from *DNMT3A*^MUT^ CH samples (left panel) and *TET2*^MUT^ CH samples (right panel) colored by the mean likelihood of cells being in the mutant clone relative the average in the HSC/MPP. As for Figure 3C-D and 3F-G, MELD was used to estimate the density of cells from each genotype within each sample using the SAM-weighted PCA as input. The mutant clone likelihood was then divided by the mean likelihood in the HSC/MPP cluster to obtain a relative likelihood, and the mean relative likelihood across all samples analyzed is shown (n = 6 *DNMT3A*^MUT^ CH samples; n = 4 *TET2*^MUT^ CH samples). A relative likelihood ≥ 1 indicates that the probability of a given cell being mutant is greater than the average for the HSC/MPP, whereas a relative likelihood ≤ 1 indicates that the probability is lower. (H) Scatterplot showing the distribution of mean mutant clone likelihoods from (G) in the HSC1 and HSC2 clusters. Each dot represents a cell, and the color scale shows the standard deviation of likelihood values across the samples. (I) Comparison of the mutant clone size between HSC1 and HSC2 clusters for *DNMT3A*^MUT^ clones (left) and *TET2*^MUT^ clones (right). Each dot represents a sample. P-values calculated by paired Wilcoxon signed-rank test. (J) GSEA against NF-κB, interferon, aging, and hematopoietic signatures comparing *DNMT3A*^MUT^ versus *DNMT3A*^WT^ HSC/MPPs (left) and *TET2*^MUT^ versus *TET2*^WT^ HSC/MPPs (right) within CH samples. Signatures with FDR > 0.2 are colored grey. Positive NES values indicate enrichment in mutant cells. * p < 0.05, ** p < 0.01, *** p < 0.001, **** p < 0.0001. (K) Model of *DNMT3A*^MUT^ and *TET2*^MUT^ clonal expansion. Exposure to inflammation impairs the function of CH^WT^ HSCs but mutant HSCs are less affected, leading to clonal expansion over time (circular arrows). Downstream of the HSC, both *DNMT3A*^MUT^ and *TET2*^MUT^ clones expand moderately in early progenitors (linear arrows). In later differentiation, *DNMT3A*^MUT^ clone size is largely maintained, but *TET2*^MUT^ clones expand further and have a myeloid bias.

Next, we asked whether the HSC clusters were transcriptionally distinct with respect to expression signatures of inflammation and aging (Figure 6C). Interestingly, HSC2 cells expressed significantly higher levels of TNFα/NF-κB signaling, aged HSC and quiescent HSC signatures,^70,90^ and lower levels of the activated HSC signature. Concordant with the more quiescent transcriptional profile, HSC2 cells expressed fewer genes than HSC1 (Figure S6D). Cluster HSC3 exhibited a more heterogeneous AUC cell score distribution and appeared transcriptionally intermediate to HSC1 and HSC2 in its transcriptional profile. This pattern of gene expression was also mirrored by expression of exemplar genes associated with inflammatory signaling, quiescence, and cell cycle across the clusters (Figures S6E-G). For example, HSC2 expressed lower levels of *CDK6*, which promotes exit from quiescence in LT-HSCs,^91^ and higher levels of *GPRC5C*, which marks dormant human BM HSCs (Figures S6F-G).^70^ Interestingly, though the HSC1 cluster showed higher expression of genes that promote exit from quiescence (Figure S6G), it also showed higher expression of genes and TF regulons implicated in HSC self-renewal (Figure S6H-I). Taken together, this suggests the HSC2 cluster has a transcriptional phenotype reflecting greater NF-κB pathway activity, increased quiescence, less proliferation, and decreased expression of transcription factors that support HSC self-renewal.

Next, we asked if these HSC clusters differed between CH and non-CH samples. To begin to address this, we first asked if the clusters differed with respect to their expression of the CH^WT^ HSC/MPP and non-CH HSC/MPP signatures (determined earlier, see Figure 5F). By AUCell score, HSC2 cells expressed significantly higher levels of the CH^WT^ HSC/MPP signature and lower levels of the non-CH HSC/MPP signature (Figure 6D). Thus, there is concordance between the elevated expression of inflammatory and aged HSC signatures in WT HSC/MPPs from CH samples and in the HSC2 cluster.

To determine if the heightened inflammatory transcriptional response in HSC/MPPs in CH versus non-CH samples correlated with differences in HSC cluster composition, we compared the frequency of the different HSC and MPP clusters as a proportion of all cells in the Lin^-^ CD34^+^ compartment (excluding Lin^-^CD34^+^CD38^-^ enriched cells) of non-CH and CH samples (Figure 6E-F). Interestingly, the HSC1 cluster was significantly smaller in CH samples. Though the HSC2 cluster was larger in CH samples, this did not reach statistical significance. Taken together, this suggests differences in HSC cluster composition between CH and non-CH samples may provide a cellular basis for the heightened response to inflammation in CH.

The analysis above did not examine differences between CH^MUT^ HSCs and WT HSCs within CH samples. Specifically, it did not provide an explanation for why the inflammatory and aging signatures were attenuated in CH^MUT^ compared to CH^WT^ HSCs. One possible hypothesis is that there is an enrichment of CH^MUT^ over CH^WT^ HSCs within the HSC1 cluster, and an over-representation of CH^WT^ HSCs in the HSC2 cluster. To examine this, we determined the ratio of CH^MUT^ to CH^WT^ HSCs in the HSC1 and HSC2 clusters (Figure 6G-H and S6J). This analysis showed that this hypothesis was incorrect. The relative density of both *DNMT3A*^MUT^ and *TET2*^MUT^ to CH^WT^ HSCs was similar across all HSC clusters. This was corroborated by lack of statistically significant difference in CH^MUT^ clone size in the HSC1 cluster compared to the HSC2 cluster (Figure 6I).

An alternative hypothesis is that the transcriptional profiles of the different HSC clusters could be differently modified by the CH mutation. To test this hypothesis, we compared enrichment of the inflammatory, aging, HSC, and progenitor signatures between *DNMT3A*^MUT^ and *DNMT3A*^WT^ HSCs in *DNMT3A*-mutant CH samples, within the three HSC clusters (Figure 6J, left panel). Specifically within the HSC2 cluster, *DNMT3A*^MUT^ cells were negatively enriched for TNFα signaling via NF-κB, aged HSC, cycle-primed HSC, and LT-HSC signatures, but positively enriched for ST-HSC, LMPP, and GMP signatures. Similar results were seen when *TET2*^MUT^ HSCs were compared to *TET2*^WT^ HSCs in *TET2*-mutant CH samples (Figure 6J, right panel). Some of these differences were shared in the HSC1 and HSC3 clusters in both *DNMT3A*^MUT^ and *TET2*^MUT^ mutant CH samples but were far less marked. These data support the hypothesis that *DNMT3A* and *TET2* mutations, either directly or indirectly, attenuate expression of transcriptional programs related to inflammatory signaling and aging, while promoting expression of programs associated with lympho-myeloid differentiation, principally in the HSC2 cluster.

## Discussion

Our study has uncovered new insights into the mechanisms whereby the human CH^MUT^ HSC population gradually gains an advantage over the vastly more numerous non-mutant HSCs.

Over time, as the inflammatory milieu of aging suppresses HSC function, the CH mutation either directly, or indirectly, attenuates the deleterious HSC response to inflammation, enabling CH^MUT^ HSCs to gain a selective advantage (Figure 6K). This insight was enabled by the first detailed single-cell examination of CH from older individuals with unperturbed hematopoiesis. These samples reflect the outcome of decades-long clonal competition, from the time of acquisition of the CH mutation to sampling, in a human bone marrow environment. Though laborious, the high-fidelity genotyping achieved with TARGET-seq+ ensures more than 90% of all bone marrow cells are genotyped, compared to ∼20% with gel bead-based approaches.^56^ Therefore, we can more confidently assign transcriptomes to either wild-type cells or CH^MUT^ clones, that usually occur at low frequency. This enables better discrimination of the transcriptional programs of CH^MUT^ and wild-type cells within the same individual. Finally, TARGET-seq+ also provides high quality scRNA-seq data, both from highly and lowly expressed genes, with better inter-cell concordance of transcript levels.

Our data show that neither *DNMT3A* nor *TET2* heterozygous mutations alter the trajectory of hematopoiesis, from HSPC to mature cells. Quantitatively, the vast majority of the steady state fitness advantage of *DNMT3A*^MUT^ and *TET2*^MUT^ clones occurs at the HSC/MPP level (Figure 6K). CH^MUT^ clones transit and differentiate normally with modest further expansion in the early progenitor compartment. In *TET2^MUT^* clones there is further 2-4-fold expansion within LMPP/GMP stages through to more mature myelomonocytic and dendritic precursor cells. Based on transcriptional kinetics across myeloid differentiation, early *TET2*^MUT^ lympho-myeloid progenitors retain less HSC and non-myeloid gene expression programs and show premature upregulation of mature programs. Our data hypothesize that this is due to increased, and premature, activity of the myeloid transcription factors CEBPA, SPI, IRF8, CEBPD, and SPIB. The ability of *TET2* mutations to promote abnormal myeloid differentiation is further highlighted by the 10-fold increased contribution of a *TET2*^-/-^ homozygous mutant clone to mature myeloid cells compared to a *TET2*^+/-^ heterozygous clone within the same individual. Finally, data from an instructive individual with both *DNMT3A*^MUT^ and *TET2*^MUT^ clones suggest that *DNMT3A*^MUT^ clones may outcompete *TET2*^MUT^ clones in the HSC/MPP compartment but that this advantage is reversed during myeloid maturation. An important caveat with this interpretation is the larger *DNMT3A*^MUT^ clone in the HSC/MPP compartment may simply reflect a clone that arose earlier in life.

Prior murine studies show that *Tet2*^-/-^,^38,49,50^ *Dnmt3a*^-/-^, and *Dnmt3a*^R878H/+^ HSPCs^47,48^ exhibit clonal advantage in inflammatory environments. Various studies have implicated exposure to different cytokines in the clonal advantage of mutant HSPCs: IL-1 through the IL-1 receptor,^50^ IL-6 through Shp/Stat3 signaling,^49^ TNFα through the TNFR1,^48^ and IFN-γ.^47^ Intestinal bacterial translocation^38^ and chronic mycobacterial infection^47^ can act as triggers for this inflammatory state. Tet2 has been shown to directly repress the pro-inflammatory cytokine IL-6 through recruitment of the histone demethylase Hdac2 and loss of Tet2 leads to elevated IL-6 levels.^38^ However, the above studies have not fully studied the differential impact of inflammation on co-existing WT and CH^MUT^ HSCs in native hematopoiesis. Specifically, these studies did not address if CH^MUT^ HSCs clonally outcompete WT HSCs despite both cell populations being adversely affected by a heightened inflammatory state, or if CH^MUT^ HSCs were more competitive because they were less impacted by the inflammatory environment. Furthermore, animal model studies may not adequately replicate human CH where clones are exposed to changing environments over decades, and where environments are heterogeneous rather than controlled.

Our data showing enrichment for transcriptional signatures of NF-κB signaling, inflammatory response and aging in both CH^WT^ and CH^MUT^ HSC/MPPs in individuals with CH compared to HSC/MPPs from age-matched individuals without CH, support the notion of a more inflammatory environment in CH. A caveat of our findings is that mean VAF of the CH clones studied by TARGET-seq+ was 14% (mean clone size 28%), which is larger than most CH clones. This may have resulted in a more inflammatory bone marrow environment than for individuals with smaller CH clones. Regardless of this caveat, within any one individual with CH that we studied, there was a reduced transcriptional impact of inflammation on CH^MUT^ HSCs compared to WT HSCs that co-exist in the same bone marrow microenvironment. Our data support two hypotheses: first, that heighted inflammation suppresses HSC function, and secondly that the CH mutation either directly, or indirectly, represses the response to inflammation within HSCs. In support of the first hypothesis are several studies showing various inflammatory stimuli, including chronic IL-1 exposure, impair HSC self-renewal and that these changes mimic those seen in aged mice.^92–94^ Furthermore, an inflammatory bone marrow niche in aged mice, including IL-1β produced by a damaged endosteum, leads to increased myelopoiesis and impaired hematopoietic recovery.^95^ In agreement with the second hypothesis are data from zebrafish where subclonal CH mutations lead to expression of pro-inflammatory genes in mature mutant myeloid cells but anti-inflammatory genes in mutant HSPCs, providing them with a relative fitness advantage.^96^ In mice when *TET2^MUT^* subclones are exposed to IL-1, they functionally outcompete WT HSCs.^50^ This is correlated with a smaller decrease in expression of genes promoting HSC self-renewal in *TET2^MUT^* compared to WT HSCs.

Further work is now needed to define the diverse drivers of inflammation in humans and the mechanisms of how human CH^MUT^ HSCs resist the detrimental effects of inflammation. The inflammatory drivers are likely to be highly heterogenous and vary over time. Despite this, there may be common final pathways of gene regulation downstream of multiple inflammatory signals. A careful examination of the human bone marrow niche in non-CH and CH individuals, with different CH clone sizes, would be helpful in this regard. This information, if combined with molecular and functional data from HSPCs and inferred rates of clonal expansion from the same individuals, may help identify the most potent putative inflammatory drivers of selection of CH^MUT^ clones over WT HSCs. These datasets, combined with functional analyses, could ultimately lead to therapies that diminish either the most important inflammatory drivers of CH^MUT^ clonal selection, or the ability of CH^MUT^ clones to resist the deleterious effect of inflammation on HSC function.

## Acknowledgements

P.V. acknowledges funding from the Medical Research Council Molecular Haematology Unit Programme Grant (MC_UU_00029/8), Blood Cancer UK Programme Continuity Grant 13008, NIHR Senior Fellowship, and the Oxford Biomedical Research Centre Haematology Theme. N.A.J. was supported by a Medical Research Council and Leukaemia UK Clinical Research Training Fellowship (MR/R002258/1). S.T. is supported by a Scatcherd European Scholarship in partnership with The Medical Research Council/Radcliffe Department of Medicine and The Clarendon Fund. M.M., R.M., B.U., M.A.S., and A.K. were funded by the Haematology Theme of the Oxford NIHR Biomedical Research Centre. Work in the laboratory of J.E.D. is supported by funds from the Princess Margaret Cancer Foundation, Ontario Institute for Cancer Research through funding provided by the Government of Ontario, Canadian Institutes for Health Research (RN380110-409786), International Development Research Centre Ottawa Canada, Canadian Cancer Society (703212), a Terry Fox New Frontiers Program project grant, University of Toronto’s Medicine by Design initiative with funding from the Canada First Research Excellence Fund, the Ontario Ministry of Health, and a Canada Research Chair. S.G.D. is funded by a Versus Arthritis Career Development Fellowship (22425). The authors thank Prof. Thomas Höfer, Dr. Verena Körber, Dr. David Cruz Hernandez, and Mr. Angus Groom for insightful comments and discussions. The authors also acknowledge the MRC WIMM Flow Cytometry Facility and Single Cell Facility. Some of the figures in this manuscript were created using BioRender.

## Author contributions

Conceptualization, N.A.J., S.T., and P.V.; Methodology, N.A.J., S.T., and A.G.X.Z., Investigation, N.A.J., S.T., B.S., M.M., M.S.N, S.S., R.M., A.K., A.G.A.N., and E.F.F.; Formal analysis, N.A.J., A.G.X.Z, S.S., G.A.G., and A.M.; Visualization, N.A.J., A.G.X.Z, and S.S.; Resources, R.M., B.U., M.A.S., S.N., B.J.L.K., A.H.T., R.A.L., R.G., A.G.A.N., and E.F.F.; Project Administration, B.W., K.W., and D.B.; Supervision, S.G.D., A.J.C., C.N., J.E.D., S.Z.X., and P.V.; Funding acquisition, N.A.J., J.E.D. and P.V.; Writing – Original Draft, N.A.J., S.T., A.G.X.Z., J.E.D., S.Z.X., and P.V.; Writing – Review & Editing, all authors.

## METHODS

### Cell culture

JURKAT human cell line was cultured in RPMI-1640 medium (Cat# 21875034, Gibco) supplemented with 10% FBS and 1% V/V Pen-Strep (Cat# 15140122, Gibco). Cells were regularly screened for Mycoplasma contamination using the MycoAlert Mycoplasma Detection Kit (Cat# LT07-218). Cells were passaged every 2-3 days and seeded at approximately 500,000 cells/mL. Cell lines were kept in a CO2 incubator at 37^°C^.

### Patient samples

Patient samples were collected from individuals undergoing elective total hip replacement (THR) surgery at the Nuffield Orthopaedic Centre, Oxford, under the Mechanisms of Age-Related Clonal Haematopoiesis (MARCH) Study. Written informed consent was obtained from all participants in accordance with the Declaration of Helsinki. This study was approved by the Yorkshire & The Humber - Bradford Leeds Research Ethics Committee (NHS REC Ref: 17/YH/0382). Exclusion criteria were: History of rheumatoid arthritis or other inflammatory arthritis, history of septic arthritis in the limb undergoing surgery, history of hematological cancer, bisphosphonate use, and oral steroid use. Patient characteristics are summarized in Table S1.

For the multi-ome analysis in young versus aged human bone marrow, bone marrow cells from young donors (26-year-old female, and 24-year-old male) were purchased from Lonza, while bone marrow samples from aged donors (70 and 77-year-old females) undergoing hip replacement surgery were collected at the Traumatology and Orthopedics Hospital Lomas Verdes (IMSS), Mexico. These elderly donors were confirmed to have no dysplasia of any hematopoietic lineages by histological and CBC analysis.^98^ Ethical approval was obtained from the Institutional Review Board (R-2012-785-092). Patient consent was obtained verbally, and as determined by the Institutional Ethical Board.

### Sample collection and processing

Trabecular bone fragments and bone marrow aspirates were obtained from the femoral canal and collected in 10 mL anticoagulated buffer containing acid-citrate-dextrose, heparin sodium and DNase. Samples of peripheral blood were collected in EDTA vacutainers.

All samples were processed within 24 hours of collection. Peripheral blood and bone marrow aspirate samples were diluted 1:1 in RPMI-1640 (Gibco) and filtered through a 70 μm cell strainer. Trabecular bone samples were manually fragmented with scissors and washed thoroughly in RPMI media with DNase to collect trabecular marrow, which was then filtered through a 70 μm cell strainer to obtain a single cell suspension and combined with the bone marrow aspirate. Mononuclear cells were then isolated by Ficoll density gradient separation (Sigma-Aldrich). For some samples, bone marrow CD34^+^ cells were purified using a CD34 MicroBead kit and MACS separation columns (Miltenyi Biotec), according to the manufacturer’s instructions. Unseparated MNCs, CD34-enriched and CD34-deplete fractions were frozen in 90% fetal bovine serum (FBS, Sigma-Aldrich) with 10% dimethyl sulfoxide (DMSO) and stored in liquid nitrogen until further use.

Peripheral blood granulocytic cell pellets isolated by Ficoll density gradient centrifugation were frozen for DNA sequencing analysis of mutations in peripheral blood.

### Targeted DNA sequencing

#### Library preparation and sequencing

Targeted DNA sequencing was performed on bone marrow MNCs and peripheral blood granulocytic DNA samples. Pre-capture DNA libraries were prepared using the KAPA HyperPlus protocol (Roche). 100 ng of genomic DNA was fragmented by enzymatic fragmentation. Following end repair and A-tailing, adapter ligation was performed using KAPA dual-indexed adapters (Roche). Library cleanup and double-sided size selection was performed using Agencourt AMPure XP beads (Beckman Coulter) to obtain fragments of ∼320 bp. Libraries were amplified by ligation-mediated PCR for 6 cycles using a KAPA HiFi HotStart high-fidelity DNA polymerase (Roche) and purified using AMPure XP beads.

Targeted capture was performed using a custom pool of biotinylated capture probes (SeqCap EZ Prime Choice, Roche) targeting 97 genes recurrently mutated in myeloid malignancies and clonal hematopoiesis spanning 347 kb (Table S2). Amplified DNA libraries were hybridized to the capture probes in pools of 10-12 samples according to the manufacturer’s instructions. The captured DNA libraries were amplified by 14 cycles of PCR using a KAPA HiFi HotStart DNA polymerase and purified using AMPure XP Beads.

Post-capture amplified DNA libraries were quantified by Qubit (Life Technologies) and size distribution and quality analyzed using a Bioanalyzer chip (Agilent Technologies). Libraries were pooled in equimolar concentrations and were sequenced on an Illumina NextSeq 500 using paired-end reads.

#### Somatic variant calling

Sequencing data were analyzed with a custom Python pipeline based on the GATK Best Practices (GATK v4.1.2.0 and Picard v2.21.3). Raw sequencing reads were converted to an unmapped BAM file and adapter sequences soft-clipped using Picard *MarkIlluminaAdapters*. Following conversion back to a FASTQ file, reads were mapped to the hg38 human reference genome assembly using the Burrows–Wheeler aligner v0.7.17^99^ with alternate contig-aware alignment. Mapped and unmapped BAM files were merged using *MergeBamAlignment* and reads from different sequencing lanes were combined. Duplicate reads were marked using Picard *MarkDuplicates* and base quality scores recalibrated with GATK BaseRecalibrator and ApplyBQSR. Somatic variant calling was performed on the pre-processed BAM files using VarDictJava v1.7.0^100^ and Mutect2^101^ in tumor-only mode. For VarDict, variants were called with a minimum variant allele frequency of 0.01, minimum base quality score of 25 and minimum supporting reads of 2, with indel realignment and removal of adapter sequences. For Mutect2, a minimum tumor LOD of 2 was used and variants were filtered for sequence context-dependent artefacts using *FilterMutectCalls* and *FilterByOrientationBias*. Indels were left-aligned and normalized using bcftools norm (v1.9). Variants were annotated using Annovar^102^. Target enrichment metrics and coverage was calculated using Picard *CollectHsMetrics* and custom scripts.

VarDict and Mutect2 variant calls were analyzed separately to identify a consensus list of high-confidence variants. The following post-processing filters were applied to VarDict calls to exclude likely sequencing artefacts:

1. Minimum of 5 variant reads for SNVs (with at least 2 reads in forward and reverse directions), or minimum of 10 variant reads for indels (with at least 4 reads in forward and reverse directions).
2. Minimum base quality score 30.
3. Minimum mapping quality score 40, except for variants in *U2AF1;U2AF1L5*, where the mapping quality was ignored. This is because in hg38, there is a duplication of the *U2AF1* gene on chromosome 21 called *U2AF1L5*, which results in reads being flagged as multi-mapped.
4. Maximum strand bias Fisher p-value of 0.0001.
5. No position bias towards beginning or end of reads.

The following post-processing filters were applied to Mutect2 calls:

1. Passed all default Mutect2 filters or only failing the *clustered_events* filter.
2. Minimum of 5 variant reads for SNVs (with at least 2 reads in forward and reverse directions), or minimum of 10 variant reads for indels (with at least 4 reads in forward and reverse directions).

Variants were flagged as likely germline, or sequencing artefacts, if any of the following applied:

1. Variant allele with a population allele frequency >1 in 1,000 according to any of three large polymorphism databases (Gnomad, 1000 Genomes Project, ESP6500) that is not a hotspot driver mutation with a COSMIC (v88) occurrence count of >100 cases or is present in a list of clonal hematopoiesis-associated mutations compiled from five _large studies._7,8,21,22,103
2. Variant allele frequency (VAF) between 0.4–0.6 or >0.9 unless recurrent in COSMIC >5 times, or previously reported in clonal hematopoiesis.
3. Present in a panel of normal cord blood samples.
4. Recurrent in the cohort unless present at least 5 times in COSMIC or at least 2 times in the clonal hematopoiesis studies.

After filtering, variants were manually inspected using the Integrative Genomics Viewer (IGV) tool (http://software.broadinstitute.org/software/igv/).

#### Annotation of pathogenic driver variants

Samples were annotated as having CH based on the presence of at least one driver mutation in bone marrow sequencing at VAF ≥ 0.01. Variants were annotated as pathogenic driver mutations using the following criteria:

1. Truncating mutations (nonsense, splice site or frameshift indel) in the following genes: *DNMT3A*, *TET2*, *ASXL1*, *NF1*, *IKZF1*, *RAD21*, *WT1*, *KMT2D*, *SH2B3*, *TP53*, *CEBPA*, *RUNX1*, *BCOR*, *KDM6A*, *STAG2*, *PHF6*, *KMT2C*, *KMT2E*, *PPM1D*, *ATRX*, *EZH2*, *CREBBP*, *NOTCH1*, *CUX1* and *ZRSR2*.
2. Non-synonymous variants at the following hotspot residues: *CBL* E366, L380, C384, C396, C404 and R420; *DNMT3A* R882; *FLT3* D835; *IDH1* R132; *IDH2* R140 and R172; *JAK2* V617F; *KIT* W557, V559 and D816; *KRAS* G12, G13, Q61 and A146; *MPL* W515; *NRAS* G12, G13 and Q61; *SF3B1* K666 and K700; *SRSF2* P95; *U2AF1* S34, R156 and Q157.
3. Non-synonymous variants occurring within the following residues of *DNMT3A*: p.292-350, p.482-614 and p.634-912; *TET2*: p.1104-1481 and p.1843-2002; or *NOTCH1*: p.1574-1620, p.1671-1721.
4. Truncating variants in *CALR* exon 9.
5. *FLT3* internal tandem duplications.
6. Non-synonymous variants reported at least 10 times in COSMIC with VAF < 0.4.
7. Non-synonymous variants falling within an annotated InterPro domain with VAF < 0.4.
8. Non-synonymous variants reported in COSMIC > 100 times.

If a variant did not meet these criteria, it was annotated as a variant of unknown significance (VUS).

#### Comparison of bone marrow and peripheral blood allele frequencies

Variants were called in peripheral blood DNA sequencing data as described above, using a minimum VAF cutoff of 0.005. Unfiltered variant calls were intersected with the list of curated bone marrow variants to compare the VAF between bone marrow and blood. For those mutations known to be present in bone marrow which were not called in peripheral blood, raw allele counts were performed directly from the BAM files using bcftools mpileup (with minimum base quality of 30 and minimum mapping quality of 35), and the allele frequency was calculated.

#### Flow cytometry and FACS sorting

Thawing media was prepared with IMDM medium (Gibco) supplemented with 20% FBS and 110 µg/mL DNase. Bone marrow samples were thawed at 37°C in a water bath, 1 mL warm FBS was added, and the suspension then diluted by dropwise addition of 8 mL thawing media. The suspension was centrifuged at 400 g for 10 mins, cells were resuspended in flow cytometry staining medium (IMDM with 10% FBS and 10 μg/mL DNase), filtered through a 35 μm cell strainer, and placed on ice.

Cells were stained with antibodies listed for 20–30 min on ice. Following antibody incubations, cells were washed with 1 mL flow cytometry staining buffer, centrifuged at 350 g for 5 min and resuspended in flow cytometry staining buffer containing the live/dead stain.

For flow cytometry analysis of BM samples, unseparated BM MNCs were used. Samples were stained with the following antibodies: anti-CD38-BV421 (1:20, Biolegend, clone HIT2), anti-CD10-BV605 (1:40, Biolegend, clone HI10a), anti-CD49f-BV650 (1:40, Biolegend, clone GoH3), anti-CD117-BV785 (1:40, Biolegend, clone 104D2), anti-CD45RA-BB515 (1:40, BD, clone HI100), anti-CD123-PE (1:40, Biolegend, clone 6H6), anti-CD90-PE/Cy7 (1:20, Biolegend, clone HI100), anti-CD34-APC (1:160, Biolegend, clone 581), anti-CD2-PE/Cy5 (1:160, Biolegend, clone RPA-2.10), anti-CD3-PE/Cy5 (1:320, Biolegend, clone HIT3a), anti-CD4-PE/Cy5 (1:160, Biolegend, clone RPA-T4), anti-CD8a-PE/Cy5 (1:320, Biolegend, clone RPA-T8), anti-CD11b-PE/Cy5 (1:160, Biolegend, clone ICRF44), anti-CD14-PE/Cy5 (1:160, eBioscience, clone 61D3), anti-CD19-PE/Cy5 (1:160, Biolegend, clone HIB19), anti-CD20-PE/Cy5 (1:160, Biolegend, clone 2H7), anti-CD56-PE/Cy5 (1:80, Biolegend, clone MEM188), anti-CD235ab-PE/Cy5 (1:320, Biolegend, clone HIR2), and propidium iodide (final concentration 3 μM; Biolegend) was used for dead cell exclusion. Analysis was done on a LSR Fortessa X20 (BD Biosciences). Unstained, single-stained (CompBeads, BD Biosciences), and fluorescence-minus-one (FMO) controls were used to determine background staining and compensation in each channel. Gating was kept consistent across all samples to enable quantification of population sizes.

For FACS sorting of BM samples for TARGET-seq+, either unseparated BM MNCs or CD34-enriched BM MNCs were used. Samples were stained with the following antibodies: anti-CD38-BV421 (1:20, Biolegend, clone HIT2), anti-CD10-BV605 (1:40, Biolegend, clone HI10a), anti-CD117-BV785 (1:40, Biolegend, clone 104D2), anti-CD45RA-BB515 (1:40, BD, clone HI100), anti-CD123-PE (1:40, Biolegend, clone 6H6), anti-CD49f-PE/Dazzle594 (1:160, Biolegend, clone GoH3), anti-CD90-PE/Cy7 (1:20, Biolegend, clone HI100), anti-CD34-APC (1:160, Biolegend, clone 581), anti-CD2-PE/Cy5 (1:160, Biolegend, clone RPA-2.10), anti-CD3-PE/Cy5 (1:320, Biolegend, clone HIT3a), anti-CD4-PE/Cy5 (1:160, Biolegend, clone RPA-T4), anti-CD8a-PE/Cy5 (1:320, Biolegend, clone RPA-T8), anti-CD11b-PE/Cy5 (1:160, Biolegend, clone ICRF44), anti-CD14-PE/Cy5 (1:160, eBioscience, clone 61D3), anti-CD19-PE/Cy5 (1:160, Biolegend, clone HIB19), anti-CD20-PE/Cy5 (1:160, Biolegend, clone 2H7), anti-CD56-PE/Cy5 (1:80, Biolegend, clone MEM188), anti-CD235ab-PE/Cy5 (1:320, Biolegend, clone HIR2), and 7-AAD (Biolegend) was used for dead cell exclusion. For sample NOC156, cells were stained with the same panel except anti-CD49f, and anti-CD117 was substituted for anti-CD117-PE/Dazzle594 (1:80, Biolegend, clone 104D2). Single-cell index sorting was performed on a Sony MA900 into 384-well plates containing 3 µL lysis buffer (except for optimization experiments, which were done in 96-well plates). Unstained, single-stained, and FMO controls were used to determine background staining and compensation in each channel. Doublets and dead cells were excluded. The following populations were sorted: Live/Lin^-^CD34^+^, Live/Lin^-^ CD34^+^CD38^-^, and CD34^-^ cells (except for sample NOC156, where only CD34^+^ cells were analyzed). In addition to the sample of interest, cells from the NOC153 control sample were sorted onto every plate, making up approximately 10% of wells on each plate, and two empty wells were used as no-template controls. After sorting, each plate was centrifuged and snap frozen on dry ice prior to storage at –80°C.

Flow cytometry data analysis was performed using FlowJo v10.8.1 and R.

### TARGET-seq+ library preparation

#### Primer design

Targeted genotyping primers used in the pre-amplification RT-PCR step were designed to amplify regions 180–900 bp long. Where possible, gDNA primers were designed to anneal within intronic regions flanking the mutation of interest, while mRNA primers were designed to anneal to exonic regions outside of the gDNA amplicon, so that independent amplicons would be generated from mRNA and gDNA. Furthermore, when a heterozygous SNP was observed close to the mutation, primers were placed in order to cover the SNP within the amplicon, enabling a direct measurement of allelic dropout. Primers were designed with Primer3Plus^104^ and specificity was checked using Primer-BLAST.^105^ For each target, primer pairs were tested for specificity and efficiency in bulk PCR reactions and in single cells.

Targeted primers for use in the genotyping PCR1 step were designed to be nested within each of the amplicons generated in the pre-amplification RT-PCR. Nested amplicons were 290– 631 bp in length. As for pre-amplification primers, gDNA primers were designed to anneal within intronic regions flanking the mutation, and cDNA primers were designed to anneal to exonic regions outside of the gDNA amplicon, to generate independent mutational readouts from cDNA and gDNA where possible. Primer pairs were tested for specificity and efficiency in bulk PCR reactions and in single cells.

#### Lysis buffer preparation

Lysis buffer was prepared as described in Table S4, consisting of 0.1% Triton X-100 (Sigma-Aldrich), 0.5 mM dNTPs (Life Technologies), 5% PEG 8000 (Sigma-Aldrich), 0.5 U/μL RNase inhibitor (Takara), 2.7 x 10^-05^ AU/μL protease (Qiagen), and 1:8,000,000 diluted ERCC spike-in mix (Ambion). 25 μL of lysis buffer was dispensed into each well of a 384-well stock plate using a Formulatrix Mantis with a high-volume chip, and 1.79 μL of 10 μM barcoded oligodT-ISPCR primer was added to each well using an INTEGRA Viaflow. 3 μL of barcoded oligodT-lysis buffer mix was then transferred into each well of a 384-well plate. Plates were sealed and stored at –80°C and thawed prior to cell sorting.

#### Reverse transcription and pre-amplification

Plates containing sorted cells were removed from –80°C storage and incubated at 72°C for 15 min to perform cell lysis, RNA denaturation and protease heat inactivation. For reverse transcription (RT), 1 µL of RT mix was added, bringing reaction concentrations to 25 mM Tris-HCl (Thermo Scientific), 30 mM NaCl (Invitrogen), 2.5 mM MgCl2 (Invitrogen), 1 mM GTP (Thermo Scientific), 8 mM Dithiothreitol (DTT, Thermo Scientific), 0.5 U/μl RNase inhibitor (Takara), 2 μM of Smart-seq2 template switching oligo (TSO, IDT), 2 U/μl of Maxima H-minus reverse transcriptase enzyme (Thermo Scientific), and target-specific mRNA primers (70 µM final concentration). RT was performed by incubation at 42°C for 90 min followed by 10 cycles of 50°C for 2 min and 42°C for 2 min. The reaction was terminated by incubating at 85°C for 5 min.

Pre-amplification PCR mix containing target-specific genotyping primers was prepared as described in Table S4, to achieve reaction concentrations of 1× KAPA HiFi HotStart Ready Mix (Roche), 50 nM ISPCR primer, 28 nM target-specific cDNA primers, and 400 nM target-specific gDNA primers. Pre-amplification PCR was performed directly after reverse transcription by addition of 6 μL PCR mix and incubation on a thermocycler using the following program: 98°C for 3 min for initial denaturation, 21 cycles of 98°C for 20 s, 67°C for 30 s and 72°C for 6 min. Final elongation was performed at 72°C for 5 min. Conditions used for all RT-PCR steps are listed in Table S4. The sequences of the primers used in the RT-PCR steps for whole transcriptome amplification and targeted genotyping amplification are listed in Table S4.

Following cDNA amplification, successful libraries contain whole transcriptome cDNA and amplicons spanning each targeted mutation. An aliquot of this cDNA-amplicon mix was used for whole transcriptome library preparation and another aliquot for single-cell genotyping library preparation. 1 μL of cDNA-amplicon mix was pooled per well to create cDNA pools from 192 uniquely barcoded single-cell libraries, using a Mosquito HTS liquid handling platform (TTP Labtech). Each cDNA pool was purified twice using Ampure XP beads with 0.6:1 beads to cDNA ratio. Pooled cDNA libraries were checked using a High Sensitivity DNA Kit on a Bioanalyzer (Agilent) or a High Sensitivity NGS Fragment Analysis Kit (1 bp - 6,000 bp) on a Fragment Analyzer (Agilent). Libraries were quantified by Qubit dsDNA HS Assay (Life Technologies). These pools were used to generate 3’ biased whole-transcriptome libraries. The remainder of the cDNA-amplicon mix was diluted 1:2 with water and stored at –20°C for use in single-cell genotyping.

#### Whole-transcriptome library preparation and sequencing

Bead-purified cDNA pools were used for tagmentation-based library preparation with a Nextera XT DNA Library Preparation Kit (Illumina) using a custom PCR amplification strategy to generate 3’ biased libraries containing oligodT cell barcodes as previously published,^60^ with some modifications. Pooled cDNA libraries were diluted to 800 pg/μL and a total of 4 ng (5 μL) from each pool was used in the tagmentation reaction with 10 μL tagmentation buffer (TD) and 5 μL ATM enzyme. The reaction was incubated at 55°C for 10 min, followed by the addition of 5 μL 0.2% SDS to release Tn5 from the DNA. Library amplification was performed using 5 μL Nextera XT i7 forward index primer (Illumina) and 5 μL custom i5 index primers (2 μM) (see Table S4 for sequences). The custom i5 index primer binds the barcoded oligodT-ISPCR adapter, resulting in amplification of the 3’ fragments containing the cell barcode. PCR was performed by adding NPM enzyme (Nextera XT DNA Library Preparation Kit, Illumina) and incubation on a thermocycler using the following program: 95°C for 30 s, 14 cycles of 95°C for 10 s, 55°C for 30 s and 72°C for 30 s, and then a final elongation of 5 min at 72°C. After tagmentation, each indexed pool was purified twice with Ampure XP beads using a 0.7:1 beads to cDNA ratio. Library quality was checked using a High Sensitivity DNA Kit on a Bioanalyzer and quantified using Qubit dsDNA HS Assay. Equimolar pools were made and sequenced using custom sequencing primers for Read1 and Index2 (P5-SEQ, I5-SEQ, 300 nM in HT1 buffer, see Table S4). For benchmarking experiments, libraries were sequenced on a NextSeq 500/550 High Output v2.5 (75 cycle) kit (Illumina) using the following sequencing configuration: 15 bp R1; 8 bp index read; 69 bp R2. For the main experiments, up to 9,984 single-cell libraries (52 pools of 192 single-cell libraries) were sequenced on a NovaSeq S4 flow cell with a targeted sequencing depth of 1 million reads/cell using the following sequencing configuration: 15 bp R1; 8 bp index read 1; 8 bp index read 2; 200 bp R2.

#### Targeted single-cell genotyping

To generate Illumina-compatible libraries for single-cell genotyping, two PCR steps were performed as previously published in the TARGET-seq protocol.^60^ As the genotyping amplicons generated by the pre-amplification RT-PCR are not barcoded, genotyping PCR reactions were carried out separately for each single-cell library.

In the first PCR step (genotyping PCR1), nested target-specific primers containing universal CS1 (forward primer) or CS2 (reverse primer) adapters are used to amplify the target regions of interest. Incorporation of a barcode sequence specific to each plate into these primers enables libraries from different plates to be pooled subsequently. Primer sequences used for genotyping PCR1 for each sample are listed in Table S4. PCR1 reactions were performed using 3.25 μL of KAPA 2G Robust HS Ready Mix (Sigma-Aldrich), 1.5 μL of diluted cDNA-amplicon mix and 300 nM target-specific primers, in a 6.5 μL reaction.

In the second PCR step (genotyping PCR2), Illumina-compatible adapters containing a 10 bp cell barcode are attached to the genotyping PCR1 product by binding to the CS1/CS2 adapters. PCR2 reactions were performed using FastStart High Fidelity polymerase (Sigma-Aldrich) with 1.0 μL of PCR1 product and 1.2 μL of each barcode primer mix (Access Array Barcode Library for Illumina Sequencers-384, Single Direction, Fluidigm) in a 6.2 μL reaction.

Indexed amplicons were pooled using a Mosquito HTS liquid handling platform and purified with Ampure XP beads using a 0.8:1 beads to PCR product ratio. Purified pools were quantified using Qubit dsDNA HS Assay and the quality checked using a Tapestation High Sensitivity D1000 kit (Agilent) to ensure the size distribution of amplicons was as expected. Each pool was diluted to a final concentration of 4 nM and further diluted to 10 pM in HT1 buffer prior sequencing. Libraries were sequenced on a NextSeq 500/550 Mid Output v2.5 kit (300 cycle) (Illumina) using 150 bp paired-end reads, with 10 bp for the cell barcode index read and custom sequencing primers (Table S4).

#### TARGET-seq+ validation experiments

Validation experiments comparing TARGET-seq+ with TARGET-seq were performed in 96-well plates using JURKAT cells and primary human CD34^+^ HSPCs. 3’ TARGET-seq libraries were generated according to the published protocol^60^. Cells were sorted into 4.1 μL lysis buffer, consisting of 0.18% Triton X-100 (Sigma-Aldrich), 1.0 mM dNTP (Life Technologies),

1.0 U/μl RNase inhibitor (Takara), 2.7 × 10−5 AU/mL protease (Qiagen), 1.0 μM barcoded oligodT-ISPCR primer. RT was performed using SMARTScribe enzyme, RNase inhibitor, Smart-seq2 TSO (1 μM final concentration) and targeted mRNA primers (700 nM final concentration). PCR pre-amplification was performed using SeqAmp DNA polymerase, ISPCR primers (50 nM final concentration) and targeted gDNA and cDNA primers. TARGET-seq+ RT-PCR was performed as described above using double volumes per well. For both conditions, 20 cycles of amplification were used for JURKAT cells and 24 cycles of amplification for HSPCs. 3’ transcriptome libraries were prepared as for TARGET-seq+ libraries detailed above and were sequenced on a NextSeq 500/550 High Output v2.5 (75 cycle) kit.

### Targeted single-cell genotyping analysis

#### Pre-processing and mutation calling

Single-cell genotyping reads were pre-processed using the custom TARGET-seq pipeline (https://github.com/albarmeira/TARGET-seq).^60^ Reads were first demultiplexed using the 384 well barcodes introduced via the genotyping PCR2 reaction, followed by demultiplexing based on plate barcodes introduced during genotyping PCR1. This generated separate fastq files for each single cell. Reads were aligned to hg38 using STAR version 2.7.3a with default settings and cDNA/gDNA amplicons were separated into different bam files, extracting reads matching the primer sequences used for targeted PCR barcoding. This allowed independent mutational information to be obtained from cDNA and gDNA amplicons. Variant calling was performed with *mpileup* (samtools version 1.1, options: --*minBQ 30*, --*count-orphans*, --*ignore overlaps*) and results were summarized using the custom pipeline.

Mutational calling on single cells was then performed with custom R scripts, separately for each mutation. Coverage for each cell was calculated as the sum of all reads across the variant locus for that cell. Empty wells routinely displayed zero or very few reads (usually up to 2), indicating no cross-well contamination. A filtering threshold was applied to remove cells where the amplicon was not detected, or where coverage was too low for reliable genotyping. The minimum coverage was 50 reads for gDNA amplicons and 30 reads for cDNA amplicons. In cells with coverage below the threshold, the amplicon was called undetected.

The single-cell variant allele frequency (scVAF) for each cell was calculated with the following formula:

#### scVAF = variant reads / coverage

PCR amplification and next-generation sequencing have an inherent error rate, which needs to be accounted for when calling cell genotypes. WT control bone marrow cells from a healthy individual sorted onto every plate enabled the error rate for each mutant allele to be determined. The scVAF threshold for calling a cell WT was set using the following formula:

#### WT scVAF threshold = mean(scVAF in WT control) + 3 × SD of scVAF in WT control

Cells with scVAF below this threshold were called WT for that amplicon. The scVAF threshold for calling a cell mutant was set using the following formula:

#### Mutant scVAF threshold = mean(scVAF in WT control) + 3 × SD of scVAF in WT control + 0.01

Cells with scVAF above this threshold were called mutant for that amplicon. Furthermore, we required a minimum number of 10 mutant reads for a cell to be called mutant. Cells with a borderline scVAF (between the WT and mutant scVAF thresholds), where the number of mutant reads was <10, or where allelic dropout (see next section) was confirmed by analysis of a germline SNP were called undetermined.

Genotyping information from gDNA and cDNA amplicons were then combined, and a consensus genotype assigned. Consensus genotypes were assigned as follows:

1. If the mutation was identified in either the gDNA or cDNA amplicon, the cell was called mutant.
2. If both amplicons were WT, the cell was called WT.
3. If the gDNA amplicon was WT but the cDNA amplicon was undetected or undetermined, the cell was called WT.
4. If the cDNA amplicon was WT but the gDNA amplicon was undetected or undetermined, the cell was called undetermined, due to the high allelic dropout rate of cDNA amplicons.

#### Allelic drop-out (ADO) estimation using germline heterozygous SNPs

Allelic dropout (ADO) rates were estimated for heterozygous germline SNPs present in the gDNA amplicon for four mutations. A scVAF threshold to detect the reference or alternate alleles was set for each SNP based on the scVAF distribution in WT control cells. Cells were called heterozygous (Het) if the VAF was within that boundary and a minimum of 10 reads were detected for each allele. Otherwise, cells were called homozygous reference (Hom Ref) or homozygous alternate (Hom Alt). SNP alleles were phased with each mutation based on the positive or negative correlation in VAF. When neither a mutation or the in-phase SNP allele were detected, a cell was assigned an undetermined genotype as we couldn’t discern whether it was WT or mutant.

For each allele, we then calculated the fraction of cells in which ADO occurred:

### ADO rate = Cells below scVAF threshold / Total number of cells

#### Inference of clonal hierarchies

In samples with multiple mutations, the pattern of mutational co-occurrence was used to determine clonal structures and assign a clonal identity to each cell as previously described.^106,107^ In samples where mutations were mutually exclusive, such as in samples

NOC131, NOC117, and NOC115, it was clear that these belonged to independent clones. In cases where mutations co-occur in the same cells, a linear or branching clonal structure may be present. We used infSCITE^108^ to determine the phylogenetic tree which represented the statistically most likely course of somatic events. As input, we used the matrix containing the mutational status for each locus in each cell and ran infSCITE with default parameters and ‘-r 200 -L 10000 -fd 0.01 -ad 0.02 -e 0.2 -p 1000’. We confirmed each phylogenetic tree was consistent with the frequency of cells of each genotype and the clonal size determined by bulk BM sequencing VAF.

The occurrence of ADO means that in some cells, a mutation that is present will not be detected. In some of these cases, we were still able to assign a cell to a clone. For example, in sample NOC002 there were 12 cells in which the *TET2* p.R1261C mutation was detected, but the ancestral *TET2* p.Q726X mutation was not detected. In these cases, we inferred that ADO of the ancestral mutation had occurred, and the cell was assigned to the appropriate daughter clone. In all cases, this was a rare occurrence consistent with our estimates of the ADO rate.

For all downstream analyses, including differential gene expression, the clone assignment rather than the raw genotype was used to categorize WT and mutant cells.

#### Analysis of FACS index data

Flow cytometry index data were recorded for each single cell during FACS sorting for TARGET-seq+. Fluorescence values were recorded for forward scatter (FSC), back-scatter (BSC; equivalent to side scatter, SSC), Lineage/live/dead, CD34, CD38, CD117, CD45RA, CD10, CD90, CD123 and CD49f (except for the NOC156 control sample). Index data were matched with single-cell identifiers based on the well coordinate and combined with genotyping calls and other metadata into a unified data set. Virtual FACS gating was performed in R based on the strategy used for sorting. Gates were set based on populations that were negative for each marker. Cells were labelled as positive or negative for each surface marker, and Boolean logic used to assign an immunophenotypic population label. For example, cells that were Lin^-^CD34^+^CD38^-^CD10^-^CD45RA^-^CD90^+^ were labelled as immunophenotypic HSC.

### Single-cell transcriptome data pre-processing

#### Mapping and transcript counting

Transcriptome sequencing data were demultiplexed into FASTQ files for each plate with a unique i7-i5 index combination using bcl2fastq. These files contained reads from up to 384 cells with shared plate indexes. A custom python pipeline was used to further demultiplex and map the sequencing reads. First, reads from each plate were demultiplexed using the 14 bp single-cell barcode sequence in Read1 using cutadapt (v3.4). Concurrently, cDNA reads (Read2) were trimmed for polyA tails, Nextera adapters and low-quality reads. This generated individual FASTQ files with single-ended cDNA reads corresponding to each single-cell barcode. Reads were then mapped to the hg38 reference genome and ERCC92 transcripts with STARsolo (v2.7.10a) using the GENCODE v38 reference gene annotation (filtered to include protein coding genes and long non-coding RNAs), and counts for each gene were obtained using default parameters except the following: ‘--soloType SmartSeq --soloFeatures GeneFull_Ex50pAS’. Sequencing and mapping quality metrics were calculated with FastQC (v0.11.9), Samtools flagstat (v1.12), MultiQC (v1.11) and the outputs of STAR.

#### Transcript detection and dropout frequency calculation

For the comparison of transcript detection sensitivity between TARGET-seq and TARGET-seq+ (Figure S1G and S1H), data were first downsampled to 5 × 10^5^ reads per cell to remove differences due to unequal sequencing depth. The number of genes detected per cell was calculated as the sum of genes with at least one assigned read.

For calculation of dropout rates (Figure S1I), data downsampled to 5 × 10^5^ reads per cell were used. A random sample of 16 cells per chemistry were compared for JURKAT and 20 cells per chemistry for HSPC. The dropout frequency for a given gene was calculated as the percentage of cells in which the gene was not detected (normalized counts < 1). Genes were divided into three groups to compare the dropout rate in: a) all expressed genes, defined as genes detected in at least 2 cells by any method; b) frequently expressed genes, defined as genes detected in >50% of all cells; and c) lowly expressed genes, defined as genes with a mean of 2–10 normalized counts per cell.

#### Cell-to-cell correlation analysis

Cell-to-cell correlations for JURKAT cells processed with each method (Figure xx) were calculated using pairwise Pearson correlations in libraries downsampled to 5 × 10^5^ reads per cell.

### Single-cell transcriptome analysis

#### Quality control, normalization, and variable gene identification

Single-cell transcriptome analysis was performed using the SingCellaR package (v1.2.1, https://github.com/supatt-lab/SingCellaR).^65^ Metadata including genotyping and FACS index data were matched with single-cell identifiers based on the plate and well coordinates. Cells meeting the following filtering criteria were included in the analysis: reads assigned to genes > 25,000; genes detected > 2,000 and < 15,000; reads assigned to ERCC transcripts < 50%; reads in mitochondrial genes < 15%. Genes expressed in fewer than 10 cells were removed.

Reads were normalized by library size using the pool normalization method with prior clustering from the scran package.^109^

#### Dimensionality reduction, data integration and clustering

Variable genes were identified by fitting a generalized linear model to the relationship between the mean expression and squared coefficient of variation (CV^2^) for the ERCC spike-ins, used to estimate technical noise (using the BrenneckeGetVariableGenes function from the M3Drop package).^110,111^ Genes for which the CV^2^ exceeded technical noise (FDR < 0.05) were considered variable, excluding mitochondrial and ribosomal genes and ERCC transcripts. This identified 17,324 variable genes which were used for principal components analysis (PCA).

Data integration was performed using Harmony^112^ to correct for sample effects, using the sample identifier as the batch, and the top 100 principal components (PCs). The top 100 Harmony-adjusted PCs were then used for Uniform Manifold Approximation and Projection (UMAP) analysis and Louvain graph-based clustering implemented in SingCellaR, with k-nearest neighbors (KNN) equal to 15. Effectiveness of the integration was confirmed by: (a) UMAP visualization pre-and post-integration, to confirm representation of all samples across cell types (Figures S2F and S3E-L); (b) comparison of cluster identities across samples, to confirm representation of all samples across clusters, while allowing for unequal distributions between samples as is expected from biological variation; (c) concordance between the cluster assignment of cells and their immunophenotype across samples. For example, we confirmed that the majority of cells in the HSC/MPP cluster consisted of immunophenotypic HSCs and MPPs in all samples, and conversely, the majority of immunophenotypic HSCs and MPPs were assigned to the HSC/MPP cluster.

Clusters were manually annotated based on gene set enrichment of published signatures, immunophenotypic surface marker expression, and expression of canonical marker genes. The SingCellaR ‘identifyGSEAPrerankedGenes’ function was used to pre-rank genes obtained from differential gene expression analysis comparing each individual cluster with all other clusters, and gene set enrichment analysis (GSEA) was performed using the fgsea package (v1.20.0) against gene sets obtained from 9 studies that have characterized human hematopoiesis.^62–70^ Marker genes differentially expressed in each cluster were identified with the SingCellaR ‘findMarkerGenes’ function, which uses a non-parametric Wilcoxon test on log-transformed, normalized counts, to compare expression levels, and Fisher’s exact test to compare the frequency of cells expressing each gene. Louvain clustering identified 28 clusters, which were collapsed into 23 main clusters based on similarity of GSEA results, marker gene expression and immunophenotype.

The HSC/MPP, LMPP, LMPP cycling, and EMPP clusters were further subclustered using the self-assembling manifolds (SAM) algorithm, using default settings with Harmony-adjusted PCs as input and using the sample identifier as the batch.^88^ The resulting SAM-weighted PCA was then used as input to generate the UMAP in Figure 6 and for Louvain clustering, which identified 7 clusters. For consistency with earlier analyses, these SAM-derived cluster assignments for LMPP, LMPP cycling, and EMPP were used throughout the paper, while cells assigned to the HSC1-3 and MPP clusters were labelled HSC/MPP in Figures 2–5.

#### Differential abundance analysis

Differential abundance between sample types (CH vs non-CH) and between mutant and WT cells within CH samples was analyzed using MELD,^97^ a single-cell compositional analysis method that quantifies the likelihood of a cellular state appearing in each sample or condition.

For the comparison between sample types (Figures 2F-G), only cells sorted as part of the total Lin^-^CD34^+^ gate were included (excluding the Lin^-^CD34^+^CD38^-^ and CD34^-^ sorting strategies), to avoid bias introduced by enrichment of CD38^-^ cells. Sample-associated densities were calculated by running MELD using the Harmony-adjusted PCs as input, and with optimal knn and beta values identified using the MELD parameter search. The mean relative density was calculated using the following formula:

#### Mean relative density = mean(Density of CH samples) / mean(Density of non-CH samples)

Thus, a mean relative density ≥ 1 indicates that the probability of observing a given cell is greater in CH samples compared to non-CH samples, whereas a relative density ≤ 1 indicates that the probability is lower in CH samples compared to non-CH samples.

To compare mutant and WT cells within CH samples (Figures 3 and 6), sample and genotype-associated densities were calculated for every genotype by running MELD using the Harmony-adjusted PCs as input, with optimal knn and beta values identified using the MELD parameter search. The relative density of single-mutant and WT cells was then calculated (using L1 normalization to enforce cell-wise sum to be 1), and normalized to the mean relative density in the HSC/MPP cluster for each sample, in order to quantify the relative expansion or contraction of the clone downstream of the HSC/MPP using the following formula:

#### Normalized likelihood = Relative likelihood of MUT:WT / mean(Relative likelihood of MUT:WT in HSC/MPP)

These values are visualized per sample in Figures 3J, 3N and S3E-L. Finally, the normalized likelihood was averaged across samples for visualization in Figures 3C-D, 3F-G, and 6G-H.

### Pseudotime analysis

Diffusion map embeddings^113^ were calculated in scanpy using the Harmony-adjusted PCs as input to the neighborhood graph, excluding the T cell, plasma cell and endothelial cell clusters.

Diffusion pseudotime was then calculated using the HSC at the extreme of the second diffusion component as the root cell. Pseudotime scores were extracted for cells in the HSC/MPP, LMPP, GMP, pDC and Monocyte clusters and plotted on the UMAP embedding to visualize the myeloid trajectory. For comparison of *TET2*^MUT^ and *TET2*^WT^ cell density along the myeloid trajectory, cells were downsampled to an equal number per sample (n = 176 cells from each of the 4 samples).

### Differential gene expression analysis

Differential expression testing was performed with a linear mixed model to account for sample covariance using the dream pipeline from the variancePartition package,^114^ which is based on limma-voom.^115^ Testing was performed on log normalized counts, using the scran normalization size factors. Genes were filtered to include only those expressed in at least 10% of cells in either group, except for pDCs and monocytes, where a 20% filter was used. A linear mixed model was fitted to each gene using ‘dream’ and differential expression testing was performed using ‘variancePartition::eBayes’. For comparisons between sample types, the sample type was used as the test variable, and the sample identifier, age, sex, and batch effects included as covariates. For comparisons between genotypes within CH samples, the clone was used as the test variable, and the sample identifier and batch effect included as mixed effect covariates. Samples were excluded from the comparison if they had less than 5 cells in either genotype, except for pDCs and monocytes where a minimum of 2 cells was used. P values were adjusted for multiple testing with Benjamani-Hochberg correction, and differentially expressed genes were defined as those with FDR < 0.1. For defining the CH^WT^ HSC/MPP and non-CH HSC/MPP signatures, thresholds of FDR < 0.1 and log2FC > 0.5 were used.

### Changes in gene expression along pseudotime

For plotting gene expression along pseudotime, log2 normalized expression data were fitted to the pseudotime rank using a generalized additive model (GAM) separately for WT and mutant cells.

### Gene set enrichment analysis (GSEA)

Gene rankings for gene set enrichment analysis (GSEA) were generated by differential gene expression testing using the dream mixed model as described above. Genes were ranked by the z statistic from dream. To perform GSEA, the fgseaMultilevel function from the fgsea package^116^ was used. Gene sets were obtained from MsigDB v7.5.1 and published studies. Hematopoietic gene sets used in GSEA and AUCell analyses relating to Figure 4-6 are listed in Table S5. Significantly enriched gene sets were filtered using the FDR as described in each figure.

### AUCell signature analysis

The AUCell package (v1.18.1)^80^ was used to quantify the gene set activity in single cells. AUCell gene-expression rankings were created using the SingCellaR ‘Build_AUCell_Rankings’ function. AUCell gene signature enrichment was then calculated using the ‘Run_AUCell’ function with the gene matrix transposed (GMT) file of gene sets. Hematopoietic gene sets were the same as those used for GSEA analysis described above. Differences in mean AUCell scores between WT and mutant cells were tested by a linear mixed model, using clone identity as the fixed effect and sample identity as mixed effects. P-values were obtained by a likelihood ratio test of the full model with the clone effect against the model without the clone effect.

### SCENIC transcription factor regulon analysis

To infer transcription factor (TF) regulon activity, regulon analysis was performed using pySCENIC.^80^ pySCENIC was run as per the workflow guidelines from Van de Sande et al.^117^ to identify candidate TF-regulons, using the filtered, pre-processed raw counts as the input, and a list of human TFs from Lambert et al.^118^. Candidate regulons were pruned using the annotations of TF motifs ‘motifs-v10nr_clust-nr.hgnc-m0.001-o0.0.tbl’, and CisTarget was applied using the ‘mc_v10_clust’ databases of known human TF motifs annotated at: a) 500 bp upstream and 100 bp downstream of the transcription start site (TSS); and b) 10 kilobases centered around the TSS. No drop-out masking was applied. Enrichment of refined TF regulons was quantified using AUCell, with default parameters. Tests for differential regulon activity were performed using a linear mixed model, as described above.

## FACS sorting and snRNA-seq for ‘in-house’ aging dataset

BM samples were thawed via slow dropwise addition of X-VIVO 10 media (LONZA) with 50% FBS and 100μg/mL DNaseI (Roche). Cells were centrifuged at 400g for 10 min, then dead cell depleted using a commercial kit (EasySep Dead Cell Removal (Annexin V) Kit, STEMCELL) per the manufacturer’s instructions. Cells were resuspended in PBS + 5% FBS and stained for 15 min at RT for fluorescence-activated cell sorting with the following antibodies: anti-CD45RA-FITC (1:50, BD, clone HI100), anti-CD90-PE (1:50, BD, clone 5E10), anti-CD19-BV711 (1:50, BD, clone SJ25C1), anti-CD49f-PE-Cy5 (1:50, BD, clone GoH3), anti-CD271-APC (1:100, Miltenyi, ME20.4-1.H4), anti-CD34-APC-Cy7 (1:200, BD, clone 581), anti-CD38-PE-Cy7 (1:200, BD, clone HB7), anti-CD10-AlexaFluor700 (1:50, BD, clone HI10a), anti-CD14-BV605 (1:200, BD, clone M5E2), anti-CD45-V500 (1:50, BD, clone HI30) and anti-CD33-BV421 (1:100, BioLegend, clone WM53). Cells were washed following staining and resuspended in PBS + 2% FBS containing propidium iodide and filtered through 40μm nylon mesh for cell sorting. Lin^-^CD34^+^CD38^-^ and Lin^-^CD34^+^CD38^+^ populations were sorted into PBS + 0.04% BSA + EDTA on a BD FACSAria Fusion or BD FACSAria III. Cells were counted and Lin^-^CD34^+^CD38^-^ and Lin^-^CD34^+^CD38^+^ cells mixed in the following manner (1:0.33 for 24yM, 1:1 for 26yF, 1:1 for 70yF, and 1:0.5 for 77yF) for downstream 10x Genomics multiome sample preparation by the Princess Margaret Genome Centre.

## Single-nucleus RNA-seq processing - In-house aging dataset

Single-nucleus RNA-seq processing was performed using Seurat 4.3.0 in R and applied to each sample before merging. The UMI count matrix (BM24M, BM26F, BM70F, and BM77F) was loaded in the R environment using Read10X. Doublets were identified using scDblFinder on the RNA with a pre-generated embedding after filtering genes expressed in more than 3 cells, cells with more than 200 features and more than 0.05 percent ribosomal genes. After filtering out doublets, quality control was further performed by filtering out cells with unique feature counts below 200 and greater than 3000, and percent mitochondrial genes above 10%. The 10x count matrix for each sample was corrected for ambient RNA contamination using SoupX and used for downstream analysis with the cells that passed quality control. The samples were merged, and the counts were normalized using Scran. The 2000 highly variable features were selected using the “vst” selection method with FindVariableFeatures in Seurat. The cells were scaled, and the samples were integrated using Harmony correcting the sample assignments as a covariate. The optimal number of Harmony-corrected PCA components for downstream analysis was assessed using an elbow plot optimizing at 10. A k-nearest neighbors graph was constructed using FindNeighbors with the Harmony corrected principal components (PCA), and clusters were identified using the louvain algorithm (resolution = 0.8). A cell-type mixed louvain cluster was sub-clustered to more effectively pull out distinct populations. UMAP on all single cells was performed using RunUMAP at 30 neighbours and 10 Harmony corrected PCA components.

## Single-cell RNA-seq processing - Ainciburu et al. dataset

The single-cell RNA-seq data was downloaded from GSE180298^86^ and processed using Seurat 4.3.0 in R and applied to each sample before merging. The UMI count matrix (young1, young2, young3, young4, young5, elderly1, elderly2, elderly3) was loaded in the R environment using Read10X. Doublets were identified using scDblFinder on the RNA after filtering genes expressed in more than 3 cells, cells with more than 200 features and more than 0.05 percent ribosomal genes. Doublets and cells with unique feature counts and percent mitochondrial genes above a sample-specific threshold were filtered out (young1: 200 > nFeature_RNA > 4000, percent.mt > 10; young2: 200 > nFeature_RNA > 2700, percent.mt > 10; young3: 200 > nFeature_RNA > 4000, percent.mt > 5; young4: 200 > nFeature_RNA >

4000, percent.mt > 5; young5: 200 > nFeature_RNA > 5000, percent.mt > 10; elderly1: 200 > nFeature_RNA > 4000, percent.mt > 10; elderly2: 200 > nFeature_RNA > 4000, percent.mt > 10; elderlyt3: 200 > nFeature_RNA > 5000, percent.mt > 10). The samples were merged and normalized using Scran. The 2000 highly variable features were selected using the “vst” selection method with FindVariableFeatures in Seurat. The cells were scaled, and the samples were integrated using Harmony correcting the sample assignments and technology (10x 3’ V2 chemistry vs 10x 3’ V3 chemistry) as covariates. The optimal number of Harmony-corrected PCA components for downstream analysis was assessed using an elbow plot optimizing at 15. A k-nearest neighbors graph was constructed using FindNeighbors with the Harmony reduction, and clusters were identified using the Louvain algorithm (resolution = 0.5). UMAP was performed using RunUMAP at 30 neighbours and 15 PCA components.

## Single-cell RNA-seq processing - Aksöz et al. dataset

This dataset consists of 10x 3’ V2 single-cell RNA-seq data from FACS-purified Lin^-^ CD34^+^CD38^-^CD90^+^CD45RA^-^ HSCs from 3 young and 3 aged donors (all male).^87^ Briefly, the raw fastq files were aligned against the GRCh38 (Ensembl 93) reference genome (10X Cell Ranger reference GRCh38 v3.1.0) and quantified using the Cell Ranger pipeline (v3.1.0) with default parameters and further processed using Seurat (v4.3.0). Quality control was performed separately for each donor by first filtering out cells with < 200 genes detected, and then retaining only cells with < 10% mitochondrial reads and gene counts that are less than double the median gene count detected in the data for that donor. Genes detected in less than 3 cells were removed. All cells that passed quality control were included in differential expression analysis.

## Aged vs Young HSC Differential Expression

Pseudobulk profiles of HSCs from each donor were created by taking the sum of all counts for each gene across cells belonging to the HSC cluster within that donor. For the in-house aging dataset, raw counts from young and aged HSC pseudobulks were modeled with DESeq and differential expression was run between aged HSC and young HSC with donor sex as a covariate. Young HSC and aged HSC-specific genes with log2FoldChange > 1 and FDR <

0.01 were retained as signatures for downstream analysis. For the Ainciburu dataset, DESeq was run on raw counts from young and aged HSC pseudobulks only within samples profiled by 10x 3’ scRNA-seq V2 chemistry to avoid technology-driven batch effects. This comparison in the Ainciburu dataset was confounded by donor sex, wherein all aged samples were male and all aged samples were female. To attenuate this, sex specific genes (X-inactivation genes XIST and TSIX, as well as ChrY genes outside of the para-autologous region) were filtered out from the DE results. Young HSC and aged HSC-specific genes with log2FoldChange > 1 and FDR < 0.01 were retained as signatures for downstream analysis. For the Aksöz dataset, raw count pseudobulks were modeled with EdgeR as implemented in the Libra (v1.0.0) package,^119^ and differential expression was run using a likelihood ratio test between aged HSC and young HSC. Gene identifiers were converted to GENCODE v38 and young HSC and aged

HSC-specific genes with log2FoldChange > 1 and FDR < 0.01 were retained as signatures for downstream analysis, excluding genes not in the GENCODE reference annotation.

The quality of each resulting signature was evaluated by scoring across donors within our CH cohort and evaluating their association with age. While we validated that aged HSC signatures from each dataset were positively correlated with age, young HSC signatures were uncorrelated with age rather than having the expected negative correlation. Thus, only aged HSC signatures were used for downstream analysis (Table S5).

## Quantification and Statistical Analysis

Data analysis and statistical tests were performed using R version 4.2.1. Plots were generated using ggplot2 (v3.3.6) or FlowJo (v10.8.1). Detail on statistical tests used in the different figures and definition of relevant summary statistics are included in each figure legend.

## RESOURCE AVAILABILITY

### Lead Contact

Further information and requests for resources and reagents should be directed to and will be fulfilled by the lead contact, Paresh Vyas.

### Materials Availability

The list of all oligo sequences designed in this study and used for single-cell genotyping can be found in Table S4. These include both target-specific oligos used in the PCR after reverse transcription, and nested barcoded target-specific oligos used in genotyping PCR1. Barcoded oligodT-ISPCR primers were kindly provided by Prof. Adam Mead and Dr. Alba Rodriguez-Meira, and the sequences are listed in Table S4.

### Data and Code Availability

Raw targeted DNA sequencing data, TARGET-seq+ scRNA-seq, and TARGET-seq+ single-cell genotyping data have been deposited at European Genome-Phenome Archive (EGA) in order to comply with ethical approvals and will be available as of the date of publication. Processed TARGET-seq+ scRNA-seq, single-cell genotyping and metadata will be made available through Figshare. Single-nucleus RNA-seq data for the in-house aged and young bone marrow dataset have been deposited in GEO.

