## Supplementary figures for "Selective advantage of mutant stem cells in clonal hematopoiesis occurs by attenuating the deleterious effects of inflammation and aging"

### SUPPLEMENTAL FIGURES AND LEGENDS

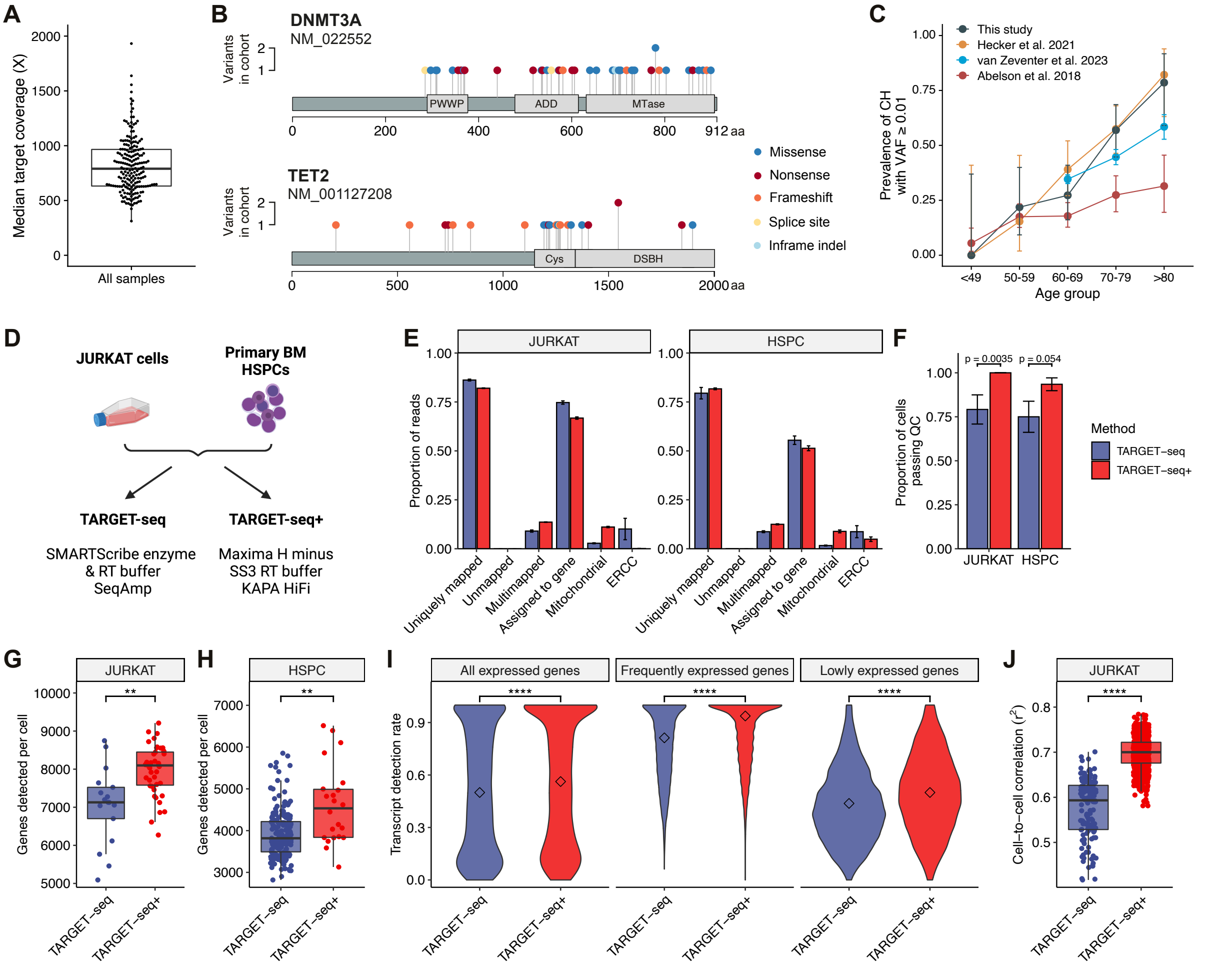


**Figure S1. Identification of cases with age-related clonal hematopoiesis in individuals undergoing hip replacement surgery and validation of TARGET-seq+, related to Figures 1-2.**

(A) Median coverage across the target region (97 genes in total) in targeted DNA sequencing of BM samples. Each point represents a sample. The boxplot shows the median and interquartile range.

(B) Distribution of mutations in coding regions of *DNMT3A* and *TET2*. Positions of protein domains are shown: PWWP, proline-tryptophan-tryptophan-proline; ADD, ATRX-DNMT3-DNMT3L; MTase, cytosine methyltransferase; Cys, cysteine-rich; DSBH, double-stranded β-helix. Mutations are colored by the predicted effect on the protein.

(C) Prevalence of CH with at least one driver mutation (VAF ≥ 0.01) by age. BM DNA sequencing data from participants in this study (n = 195) are compared with a cohort of 676 individuals reported by Abelson et al.[22], a cohort of 3,359 individuals reported by van Zeventer et al.[58] and a cohort of 199 individuals undergoing total hip replacement (n = 109 BM samples and n = 91 PB samples) reported by Hecker et al.[59] Error bars represent 95% confidence intervals.

(D) Experimental scheme for validation of TARGET-seq+ in JURKAT cells and primary human $CD34^+$ HSPCs. Metrics were compared to single cell transcriptome libraries generated with the original TARGET-seq protocol.

(E) Sequencing statistics of single cell transcriptome libraries from JURKAT and primary $CD34^+$ HSPCs processed with TARGET-seq and TARGET-seq+. Bars represent the proportion of reads for each statistic (labelled below) and error bars represent the standard error of the mean.

(F) Proportion of cells passing quality control (QC) for transcriptome libraries, for each method (n = 24 JURKAT for TARGET-seq; 46 JURKAT for TARGET-seq+; 24 HSPC for TARGET-seq; and 46 HSPC for TARGET-seq+). Error bars represent the standard error of the mean. P-values calculated by Fisher's exact test.

(G and H) Number of genes detected per cell in JURKAT cells (G) and primary $CD34^+$ HSPCs (H). In both cases, reads were downsampled to $5 \times 10^5$ reads per cell. Each dot represents a cell and each boxplot represents the median and first and third quartiles. P-values calculated by unpaired two-tailed *t*–test. ** $p < 0.01$.

(I) Comparison of transcript detection rates between TARGET-seq and TARGET-seq+ for JURKAT cells. A subsample of 16 cells per chemistry was analyzed. Reads were downsampled to $5 \times 10^5$ reads per cell. Dropout frequencies are shown for all genes expressed in at least 2 cells (left), for genes expressed in >50% of cells by any method (middle), and for genes with low expression (mean normalized counts of 2-10 per cell; right). Diamonds show the median detection rate for each condition. P-values calculated by two-tailed Wilcoxon rank-sum test.

(J) Quantification of the reproducibility of gene expression in JURKAT cells, for TARGET-seq (n = 17 cells) and TARGET-seq+ (n = 43 cells). $r^2$ values for all pairwise cell-to-cell Pearson's correlations in libraries downsampled to $5 \times 10^5$ reads per cell are shown. P-values calculated by unpaired two-tailed *t*–test. **** $p < 0.0001$.

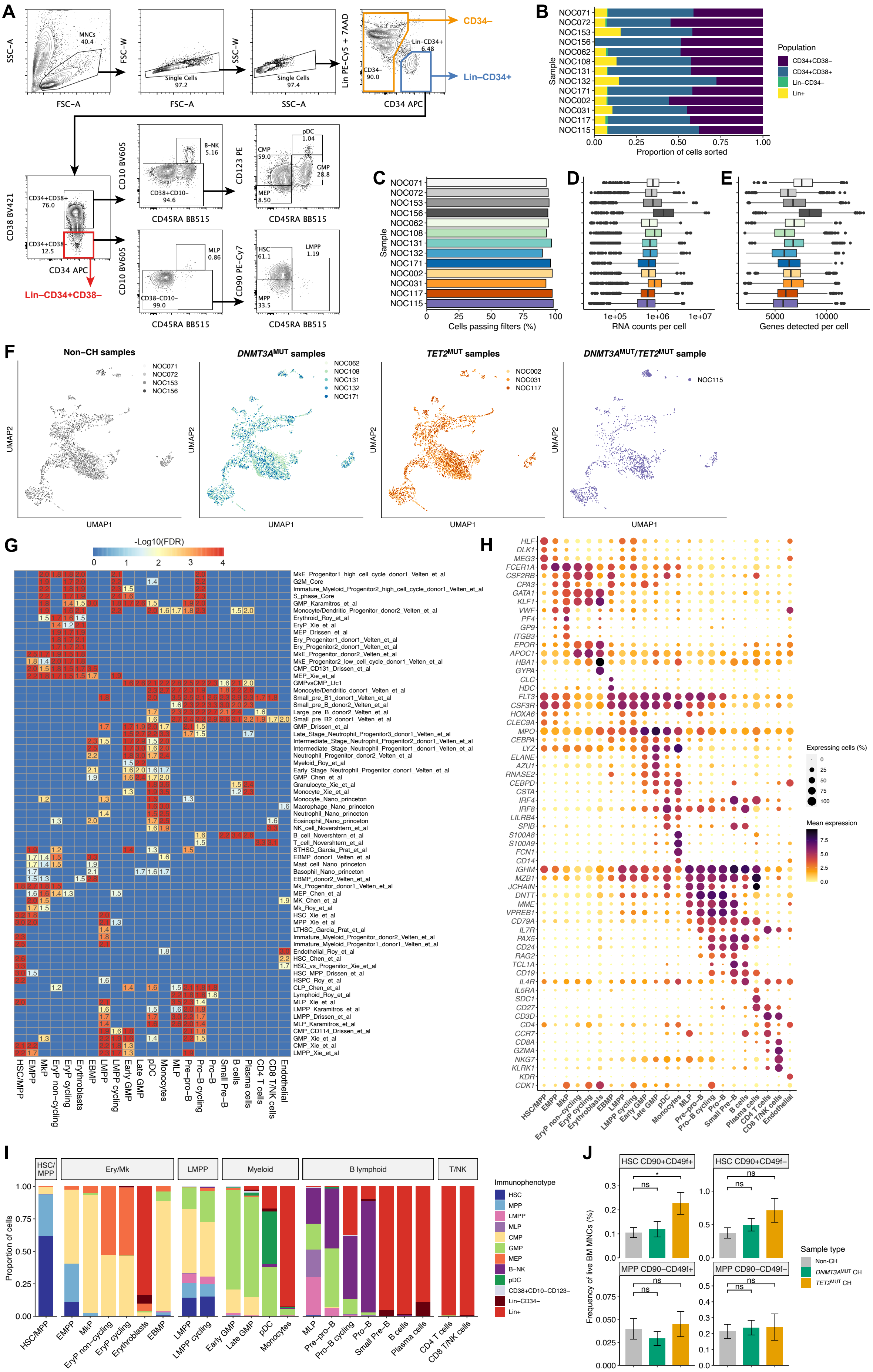
A
SSC-A
FSC-A
MNCs 40.4
FSC-W
Single Cells 97.2
SSC-W
Single Cells 97.4
Lin PE-Cy5 + 7AAD
CD34 APC
CD34– 90.0
Lin-CD34+ 6.48
CD34–
Lin–CD34+
CD38 BV421
CD34+CD38+ 76.0
CD34+CD38– 12.5
Lin–CD34+CD38–
CD10 BV605
CD45RA BB515
B-NK 5.16
CD38+CD10– 94.6
CD123 PE
CMP 59.0
pDC 1.04
GMP 28.8
MEP 8.50
MLP 0.86
CD38–CD10– 99.0
CD90 PE-Cy7
HSC 61.1
LMPP 1.19
MPP 33.5
B
Sample
NOC071
NOC072
NOC153
NOC156
NOC062
NOC108
NOC131
NOC132
NOC171
NOC002
NOC031
NOC117
NOC115
Proportion of cells sorted
Population
CD34+CD38–
CD34+CD38+
Lin–CD34–
Lin+
C
Cells passing filters (%)
D
RNA counts per cell
E
Genes detected per cell
F
Non–CH samples
DNMT3AMUT samples
TET2MUT samples
DNMT3AMUT/TET2MUT sample
UMAP1
UMAP2
G
-Log10(FDR)
H
Expressing cells (%)
Mean expression
I
Proportion of cells
HSC/MPP
Ery/Mk
LMPP
Myeloid
B lymphoid
T/NK
Immunophenotype
HSC
MPP
LMPP
MLP
CMP
GMP
MEP
B–NK
pDC
CD38+CD10–CD123–
Lin–CD34–
Lin+
J
HSC CD90+CD49f+
HSC CD90+CD49f–
MPP CD90–CD49f+
MPP CD90–CD49f–
Frequency of live BM MNCs (%)
ns
Sample type
Non-CH
DNMT3AMUT CH
TET2MUT CH

**Figure S2. TARGET-seq+ applied to *DNMT3A* and *TET2*-mutant clonal hematopoiesis, related to Figure 2.**

(A) FACS index sorting strategy for TARGET-seq+ experiments. Sorting gates for $Lin^{-}CD34^{+}$ (blue), $Lin^{-}CD34^{+}CD38^{-}$ (red), and $CD34^{-}$ (orange) cells are shown.

(B) Proportion of cells obtained in each immunophenotype for TARGET-seq+ analysis in each sample.

Panels (C) to (E) show transcriptome quality control and sequencing statistics by sample. In panels (D) and (E), boxplots show the median and interquartile range.

(C) Percentage of cells passing quality control filters.

(D) Sequencing depth (RNA counts) per cell.

(E) Number of genes detected per cell.

(F) UMAPs colored by sample, split by sample genotype. Colors indicate cells from different samples.

(G) Heatmap of GSEA results for genes differentially expressed between each cluster and all other clusters, using published gene signatures for HSPCs.[62-70] Heatmap colors represent –$\log_{10}$ of the false discovery rate (FDR). Numbers inside the heatmap indicate the normalized enrichment score for each comparison, showing only those with FDR < 0.05.

(H) Expression of marker genes by cluster. The size of each dot represents the percentage of cells expressing each gene and the color represents the mean expression level (log normalized counts).

(I) Proportions of each immunophenotypic population in each transcriptional cluster.

(J) Barplots showing the frequency of $CD90^{+/-}$ and $CD49f^{+/-}$ HSC/MPP cells as a percentage of total BM MNCs. Data are represented as mean ± SEM. P-values calculated by Wilcoxon rank sum test with Holm-Bonferroni multiple testing correction. * $p < 0.05$.

HSC, hematopoietic stem cell; MPP, multipotent progenitor; EMPP, erythroid/megakaryocyte-primed multipotent progenitor; MkP, megakaryocytic progenitor; EryP, erythroid progenitor; EBMP, Eosinophil-basophil-mast cell progenitor; LMPP, lymphoid-primed multipotent progenitor; GMP, granulocyte-monocyte progenitor; pDC, plasmacytoid dendritic cell progenitor; MLP, multi-lymphoid progenitor; B-NK, B and NK cell progenitor.

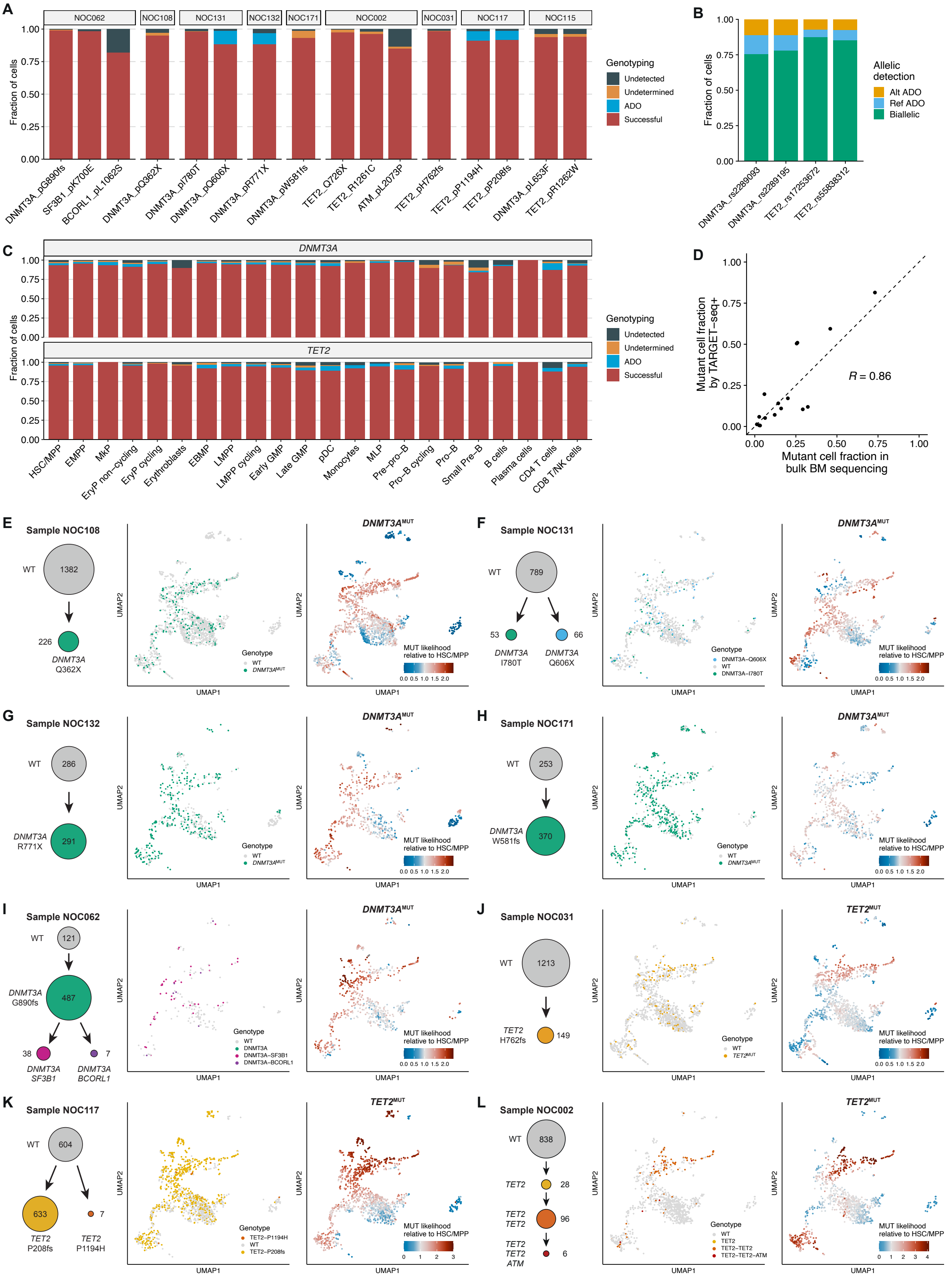

**Figure S3. TARGET-seq+ single-cell genotyping analysis, related to Figure 3.**

(A) Fraction of cells successfully genotyped (red) for each mutation (below) and sample (above) analyzed. In 4 amplicons, allelic dropout (ADO) of the mutant allele (blue) could be identified by analysis of a germline heterozygous SNP in the same amplicon. Cells with borderline VAF were assigned an undetermined genotype (orange) for that mutation.

(B) Frequency of biallelic and mono-allelic detection of 4 germline heterozygous SNPs that were co-amplified with somatic mutations.

(C) Fraction of cells successfully genotyped for *DNMT3A* and *TET2* across cell types (below). Labels are as in (A).

(D) Comparison of mutant cell fraction for each mutation identified in single cells by TARGET-seq+ (y-axis) with the clonal cell fraction as measured by bulk BM DNA sequencing (x-axis). R indicates the Pearson correlation coefficient.

(E) to (L) Clonal structures and patterns of clonal expansion for individual *DNMT3A* and *TET2*-mutant CH samples. For each sample, the clonal structure determined by single-cell genotyping is shown on the left. Number of cells assigned to each genotype are shown. The middle panels show successfully genotyped cells from each sample on the UMAP embedding, labelled by cell genotype. Right-hand panels show the relative likelihood for mutant clones carrying a single mutation in *DNMT3A* or *TET2* on the UMAP. In each case, the mutant clone likelihood is normalized to the mean likelihood in the HSC/MPP cluster. Values > 1 indicate clonal expansion relative to HSC/MPP and values < 1 indicate smaller clone size relative to HSC/MPP.

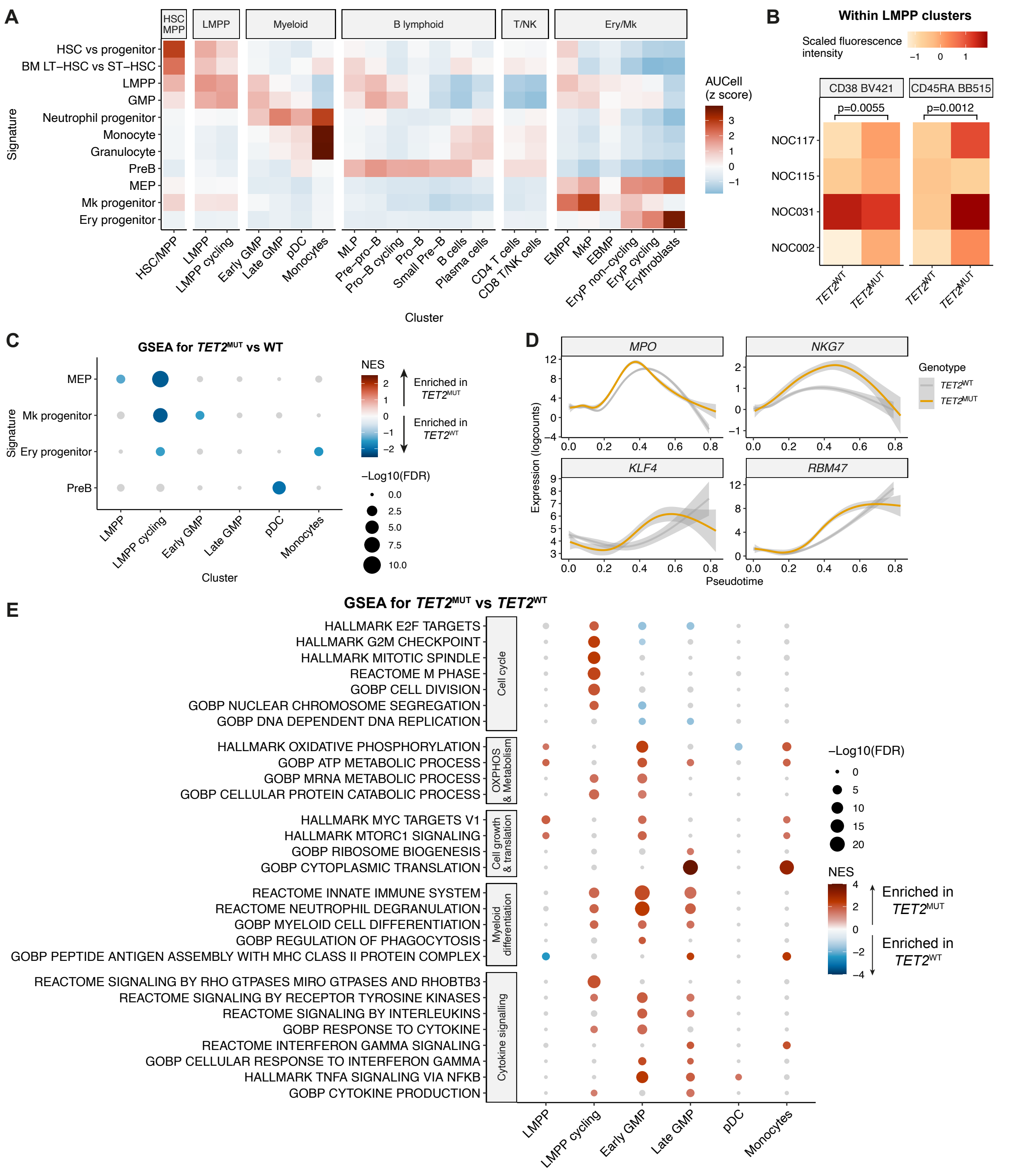


**Figure S4. Consequences of *TET2* mutations in myeloid progenitors, related to Figure 4.**

(A) Heatmap of scaled AUCell expression scores for HSC, myeloid, B-lymphoid, megakaryocytic, and erythroid gene signatures (y-axis) used in Figures 4C and S4C, by cluster (x-axis).

(B) Expression of CD38 and CD45RA cell surface protein, measured by FACS indexing in *TET2*$^{WT}$ and *TET2*$^{MUT}$ cells, in the LMPP clusters. P-values calculated by linear mixed model.

(C) GSEA against erythroid, megakaryocytic, and lymphoid gene signatures, comparing *TET2*$^{MUT}$ against *TET2*$^{WT}$ cells within each LMPP and GMP cluster. Differential expression analysis was performed accounting for sample and batch effects. Cells from the 4 *TET2*$^{MUT}$ CH samples were included. Color intensity indicates the normalized enrichment score (NES); Positive NES values indicate enrichment in mutant cells. Signatures with FDR > 0.1 are colored grey.

(D) Local regression of gene expression values along myeloid pseudotime for myeloid lineage-affiliated genes, comparing *TET2*$^{MUT}$ and *TET2*$^{WT}$ cells.

(E) GSEA against Hallmark, Gene Ontology biological process (GOBP) and Reactome signatures comparing *TET2*$^{MUT}$ against *TET2*$^{WT}$ cells within the LMPP, GMP, pDC and monocyte clusters. Cells from the 4 *TET2*$^{MUT}$ CH samples were included in the analysis.

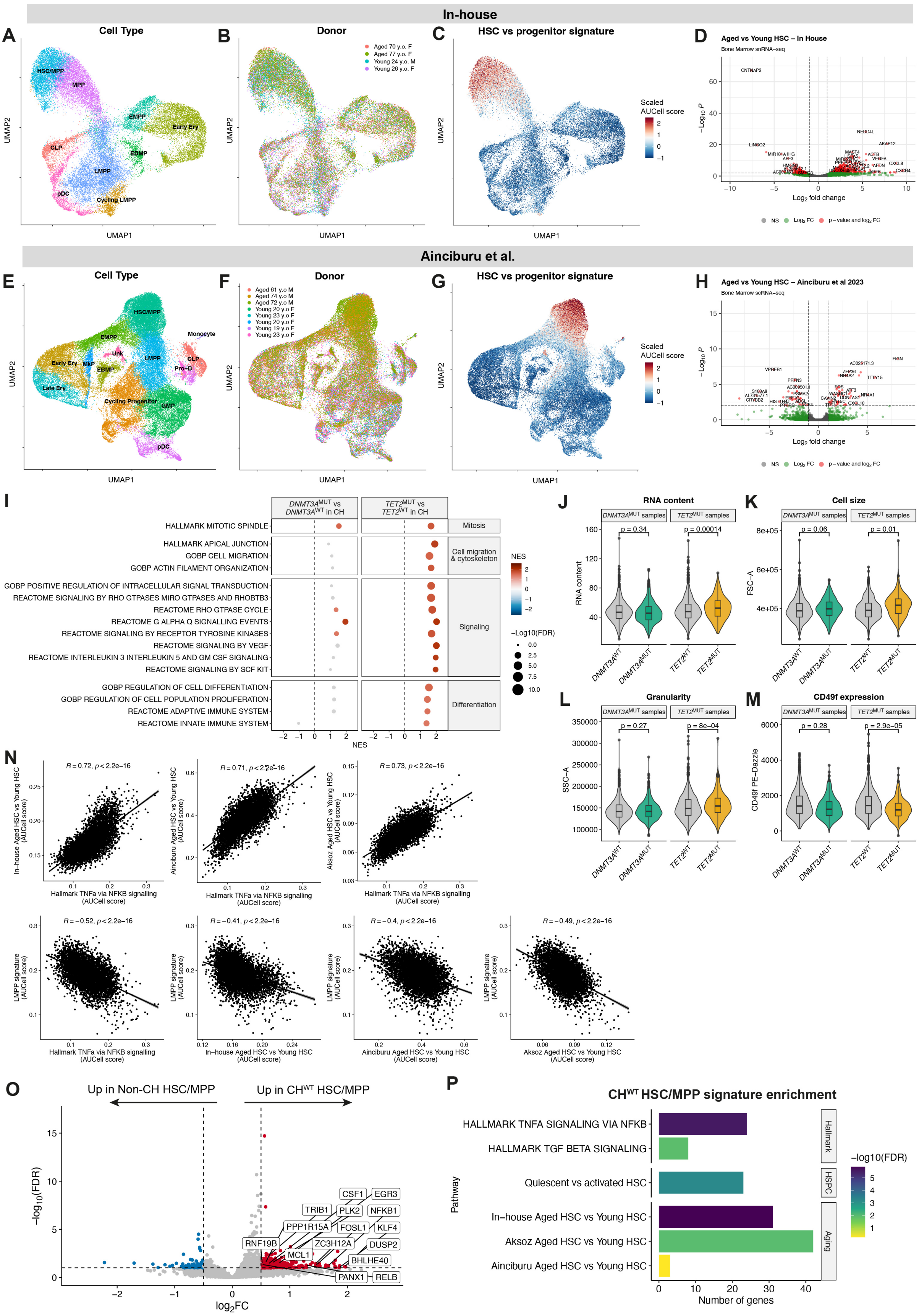


**Figure S5. Generation of aged HSC gene signatures and consequences of *DNMT3A* and *TET2* mutations in HSC/MPP, related to Figure 5.**

(A) to (C) UMAPs showing snRNA-seq data for BM HSPCs from 2 young and 2 aged donors (in-house dataset) colored by Louvain cluster identity (A), donor identity (B) and the scaled AUCell score for the HSC vs progenitor signature (C).

(D) Volcano plot showing differentially expressed genes (DEGs) between HSC/MPP from aged donors versus young donors from the in-house dataset. Gene with FDR < 0.01 and $log_2FC > 1$ were considered differentially expressed (labelled red).

(E) to (G) UMAPs showing snRNA-seq data for BM HSPCs from 5 young and 3 aged donors (Ainciburu et al. dataset[86]) colored by Louvain cluster identity (E), donor identity (F) and the scaled AUCell score for the HSC vs progenitor signature (G).

(H) Volcano plot showing differentially expressed genes (DEGs) between HSC/MPP from aged donors versus young donors from the dataset from Ainciburu et al.[86] Gene with FDR < 0.01 and $log_2FC > 1$ were considered differentially expressed (labelled red).

(I) GSEA against Hallmark, Gene Ontology biological process (GOBP), and Reactome signatures, comparing *DNMT3A*$^{MUT}$ against *DNMT3A*$^{WT}$ HSC/MPPs (left) and *TET2*$^{MUT}$ against *TET2*$^{WT}$ HSC/MPPs (right), within CH samples. Signatures with FDR > 0.2 are colored grey. Positive NES values indicate enrichment in mutant cells.

(J) RNA content (ratio between endogenous RNA reads and ERCC spike-in reads) comparing *DNMT3A*$^{WT}$ against *DNMT3A*$^{MUT}$ HSC/MPPs and *TET2*$^{WT}$ against *TET2*$^{MUT}$ HSC/MPPs. P-values calculated by linear mixed model.

(K) to (M) Cell size (K), granularity (L), and CD49f protein expression (M) measured by FACS indexing, comparing *DNMT3A*$^{WT}$ against *DNMT3A*$^{MUT}$ HSC/MPP and *TET2*$^{WT}$ against *TET2*$^{MUT}$ HSC/MPP. P-values calculated by linear mixed model.

(N) Top row: Correlation between AUCell scores for the Hallmark TNFα via NF-κB signature and aged HSC signatures within the HSC/MPP cluster. Bottom row: Correlation between AUCell scores for the LMPP signature and the Hallmark TNFα via NF-κB and aged HSC signatures within the HSC/MPP cluster (n = 4951 cells from 13 samples). For each comparison, the Spearman correlation coefficient, the associated p-value, and regression line are shown.

(O) Volcano plot showing differentially expressed genes (DEGs) between CH WT HSC/MPPs from 9 CH samples and WT HSC/MPPs from 4 non-CH samples (n = 1280 CH WT cells, 2362 non-CH cells). Genes with FDR < 0.1 and $log_2FC > 0.5$ were considered differentially expressed. Genes in the TNFα via NFKB pathway are labelled.

(P) Pathway enrichment of genes upregulated in CH WT HSC/MPPs against Hallmark and hematopoietic signatures. Hypergeometric test.

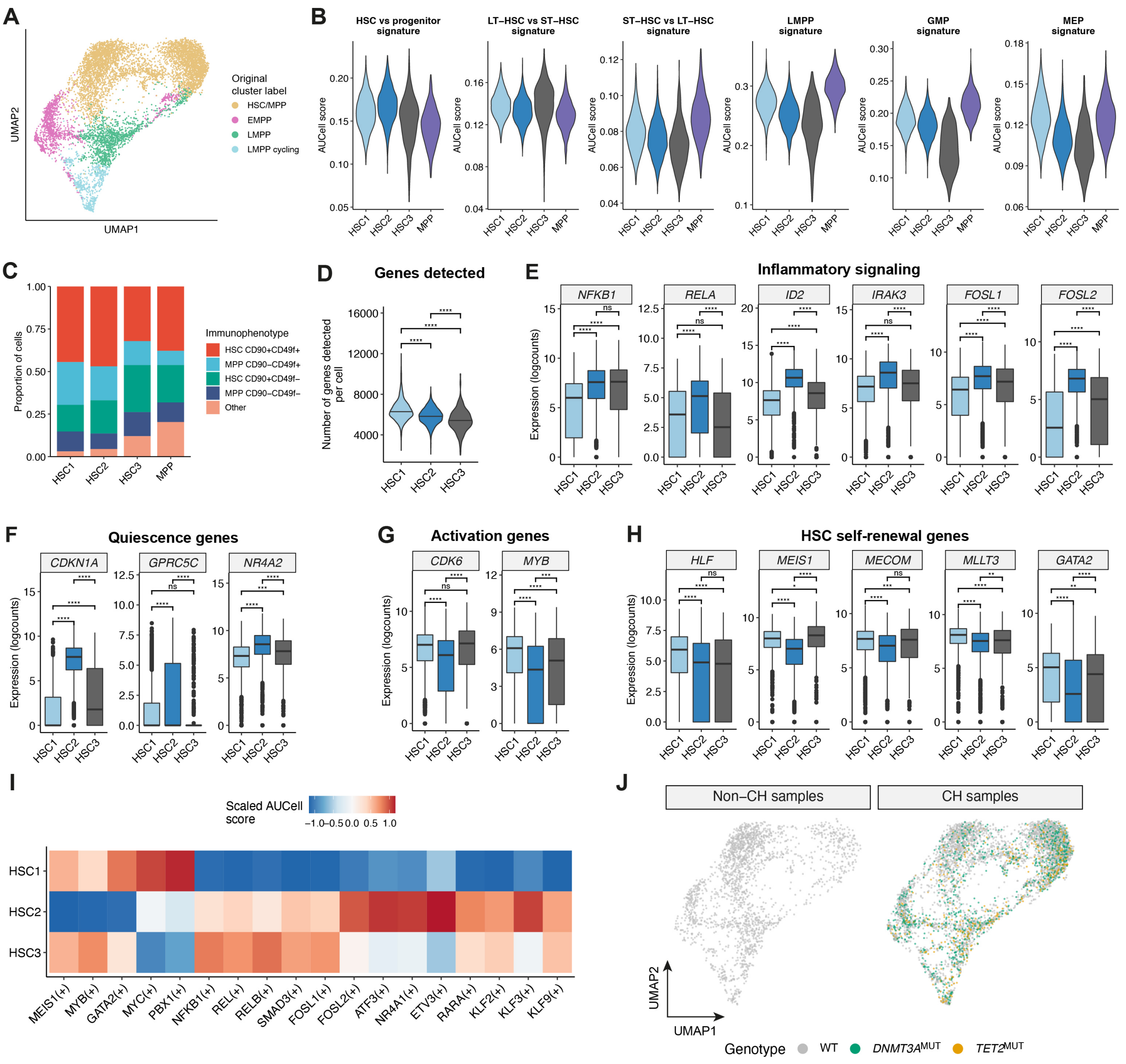


**Figure S6. Characterization of transcriptional heterogeneity within HSCs, related to Figure 6.**

(A) UMAP embedding of 8059 cells from the HSC/MPP, EMPP, LMPP and LMPP cycling clusters after feature weight derivation with the Self-Assembling Manifolds (SAM) algorithm. Cells are colored by the original cluster annotation used in previous figures.

(B) Comparison of AUCell expression scores for HSC and progenitor gene signatures (indicated above each graph) in HSC and MPP subclusters (x-axis).

(C) Immunophenotype of cells (colors) in HSC subclusters (x-axis).

(D) Violin plots showing the number of genes detected per cell within the HSC1-3 clusters. **** $p < 0.0001$. P-values calculated by pairwise *t* test.

(E) Expression of genes related to inflammatory signaling within the HSC1-3 clusters. Asterisks represent FDR-corrected p-values from differential expression testing. * FDR < 0.05, ** FDR < 0.01, *** FDR < 0.001, **** FDR < 0.0001.

(F) As in (D) but for genes related HSC quiescence.

(G) As in (D) but for genes related HSC activation.

(H) As in (D) but for genes related HSC self-renewal.

(I) Heatmap showing scaled AUCell scores for selected regulons that are differentially active in the HSC1-3 clusters.

(J) UMAP embeddings showing cells from non-CH and CH samples. Cells are colored by genotype. For clarity, only WT and single-mutant cells are shown.
